# Adaptive behaviors emerge from Hebbian-like plasticity through flexible attractors

**DOI:** 10.64898/2026.11.18.767638

**Authors:** Chenguang Li, Jonah Brenner, Hanming Ye, Benjamin Du Pont, Matas Kudarauskas, Joanna Aloor, Chunyu A. Duan, Jason Chen, Jeffrey C. Erlich

## Abstract

Adaptive behavior is commonly framed as an optimization problem, in which organisms learn to maximize reward. Here we show that simple networks with local, Hebbian-like learning rules, which have no reward objective, nonetheless reproduce hallmark features of animal reward-seeking when placed in closed-loop environments where rewards arrive as ordinary sensory input. Six canonical local learning rules were tested under these conditions, and despite most rules being designed for unsupervised representation learning, all rules learned to seek out signal. However, at the parameters tested, only one rule, prospective configuration, robustly captured all of a set of tested phenomena: reward-magnitude-dependent learning rates, probability matching, memory retention across contexts, and switching between exploration and exploitation with heavy-tailed statistics. In these networks, actions that produce input become self-sustaining states (attractors) of network dynamics. Attractors reside in an activity-weight space and mediate behaviors, and adapt to their environments rather than being fixed properties of the networks. These flexible geometries provide a parsimonious account of how heavy-tailed runs of repeated choices, which resemble Lévy flights seen in foraging, also change with the environment. A single framework then makes predictions about behavior and neural population geometries that can be used to investigate the learning dynamics of the brain.

## 1 Introduction

Inside an animal’s nervous system, individual neurons learn from each other’s activities and somehow produce the behavior of a single, coordinated organism. It is often thought that this degree of coordination requires signals that are broadcast throughout the nervous system, telling neurons what to learn, and when. ^1–3^ However, this raises the question of where such signals originate in the brain, and how they adapt to and interact with responsive environments. And while such broadcasted signals do exist and guide goal-directed behavior, much less is understood about whether plasticity driven by local neural activity can achieve goal-directed behavior on its own.

Many local learning rules have been proposed since the advent of Hebbian learning, ^4,5^ and most were designed for representation learning or memory tasks ^6–8^. Such rules did not distinguish reward from other signals, and because of this, seemed unable to support goal-driven behavior. Scientists then considered ways to deliver reward information as additional explicitly defined terms, ^9^ and this idea underlies the successes of deep reinforcement learning, ^10,11^ foundational predictions in neuroscience, ^1,12^ and the framework of three-factor rules, which add neuromodulation into the Hebbian template. ^13–15^ Today, three-factor frameworks still tend to assume, explicitly or implicitly, that plasticity based on pre- and postsynaptic activity alone (i.e. two-factor learning) cannot learn reward-seeking tasks. ^13,16^ There are exceptions: matching emerges from plasticity driven by the covariance between reward and neural activity ^17^ and local rules have been designed for closed-loop control and foraging ^18–20^. However, these rules either include a reward term or were built for a specific task. Whether canonical rules designed for other purposes can learn to broadly seek reward has not been systematically tested.

Here we revisit these assumptions. We consider that the rewards animals receive can themselves be sensory events, or be cued by them ^21,22^—the taste of sugar, or the sound of a bell—and that these signals reach neurons as ordinary inputs. Animals will also work for sensory stimulation alone, ^23–25^ and some forms of reward learning survive the loss of dopamine. ^26^ We therefore deliver reward to networks as ordinary input, allowing reward to shape learning only through activity propagated through the network. All neural activity is treated the same way, regardless of whether it is a direct response to environmental stimuli or passed through hidden layers.

We designed a unified framework to test six canonical local learning rules across a diverse suite of tasks and compare them to a three-factor rule that broadcasts a reward-prediction error ^13^ (Table 1). In this framework, behaviors emerge fully online and autonomously in closed-loop environments, without direct optimization on a global target. The tested rules are Bienenstock-Cooper-Munro (BCM), ^27^ Oja’s rule, ^28^ Law and Cooper, ^29^ prospective configuration, ^30^ covariance-based plasticity,^17^ and latent predictive learning ^31^. All were capable of signal-seeking behavior in simple leaky integrator networks. In addition, several rules parsimoniously account for long-studied behavioral phenomena, including reward-magnitude-based learning rates, ^32^ probability matching, ^33–35^ costly stochasticity,^36^ discrete explore/exploit dynamics, ^37–40^ heavy-tailed foraging distributions, ^41,42^ interference-driven forgetting, ^43,44^ and memory retention across contexts. ^45^ At the parameters tested, a single rule, prospective configuration, both performed well and was analytically tractable within our framework.

**Table 1:**
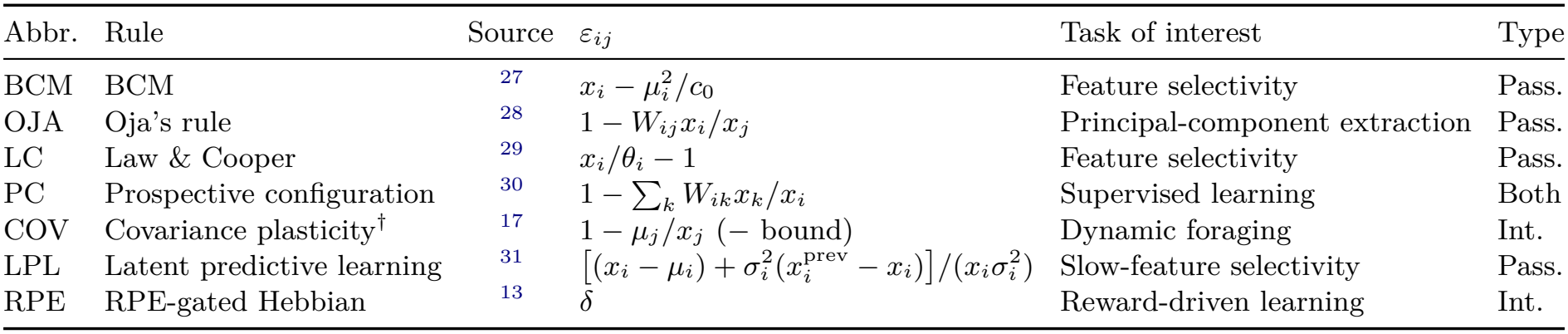
The seven plasticity rules. Rules fit a common template (Eq. 1) and differ only in *ε*_*ij*_ (*j* presynaptic, *i* postsynaptic). The running statistics of activity are the mean *µ*, the mean square *θ*_*i*_ and the variance 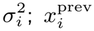 is activity on the previous timestep. The input-layer reward-prediction error 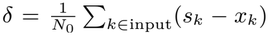. The BCM rule includes a constant *c*_0_ = 0.1. “Task of interest” is the problem each rule was originally designed for; “Type” is interactive (Int.) or passive (Pass.). ^†^COV was originally proposed with a reward term, in which plasticity follows the covariance between reward and neural activity. We use an unmodulated two-factor form, so reward reaches the network only as sensory input (Methods).

| Abbr. | Rule | Source | ε<sub>ij</sub> | Task of interest | Type |
|---|---|---|---|---|---|
| BCM | BCM | <sup>27</sup> | x<sub>i</sub> − μ<sub>i</sub><sup>2</sup>/c<sub>0</sub> | Feature selectivity | Pass. |
| OJA | Oja’s rule | <sup>28</sup> | 1 − W<sub>ij</sub>x<sub>i</sub>/x<sub>j</sub> | Principal-component extraction | Pass. |
| LC | Law & Cooper | <sup>29</sup> | x<sub>i</sub>/θ<sub>i</sub> − 1 | Feature selectivity | Pass. |
| PC | Prospective configuration | <sup>30</sup> | 1 − ∑<sub>k</sub> W<sub>ik</sub>x<sub>k</sub>/x<sub>i</sub> | Supervised learning | Both |
| COV | Covariance plasticity<sup>†</sup> | <sup>17</sup> | 1 − μ<sub>j</sub>/x<sub>j</sub> (− bound) | Dynamic foraging | Int. |
| LPL | Latent predictive learning | <sup>31</sup> | [(x<sub>i</sub> − μ<sub>i</sub>) + σ<sub>i</sub><sup>2</sup>(x<sub>i</sub><sup>prev</sup> − x<sub>i</sub>)]/(x<sub>i</sub>σ<sub>i</sub><sup>2</sup>) | Slow-feature selectivity | Pass. |
| RPE | RPE-gated Hebbian | <sup>13</sup> | δ | Reward-driven learning | Int. |

We investigate mechanisms behind the behaviors, focusing on the prospective configuration rule, ^30,46,47^ and find that behaviors come from attractor geometries that emerge from the interactions of networks and their environments. This work differs from earlier local rules for reward-seeking in two ways: in generality, as canonical rules built for other purposes all learn to seek signal, and in mechanism, as the attractors that mediate behavior are not specified in advance but emerge and change with the environment. Local plasticity may thus provide an adaptive substrate for neuromodulation to act upon, rather than neuromodulation being the sole driver of reward-seeking behavior. Overall, this work points to a surprisingly simple way for goal-directed behavior to emerge from networks of interacting neurons.

## 2 Results

### 2.1 A behavioral framework for local Hebbian-like plasticity

Six local plasticity rules and a three-factor reward prediction error (RPE)-modulated rule were compared within one framework (Table 1). All seven rules fit a common template:

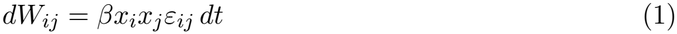

Here *W*_*ij*_ is the weight from presynaptic neuron *j* to postsynaptic neuron *i*; *β* is the weight learning rate, *x*_*i*_ and *x*_*j*_ are the activity of the post- and pre-synaptic neurons, and *ε*_*ij*_ is a rule-specific error term. For all but the three-factor rule (“RPE” in Table 1), *ε*_*ij*_ depends on local quantities.

This unification lets us compare all seven rules in both representation learning and adaptive behavior tasks (Fig. S1–S3). In all cases, neural activities follow noisy leaky integrator dynamics:

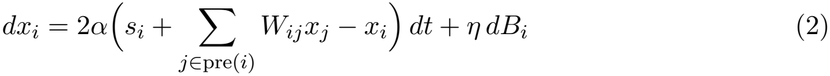

For input neurons, activities track the sensory input. For all other neurons, they track presynaptic input. In Eq. 2, *s*_*i*_ is the sensory input to neuron *i* (0 for non-input neurons), *α* is the activity update rate, *η* is the noise scale, and *B*_*i*_ is an independent standard Wiener process. Activities are rectified at zero after each update (Methods). Altogether, weights, activities, and a network’s environment determine a dynamical trajectory through the network’s state space (Fig. 1a–c).

**Figure 1:**
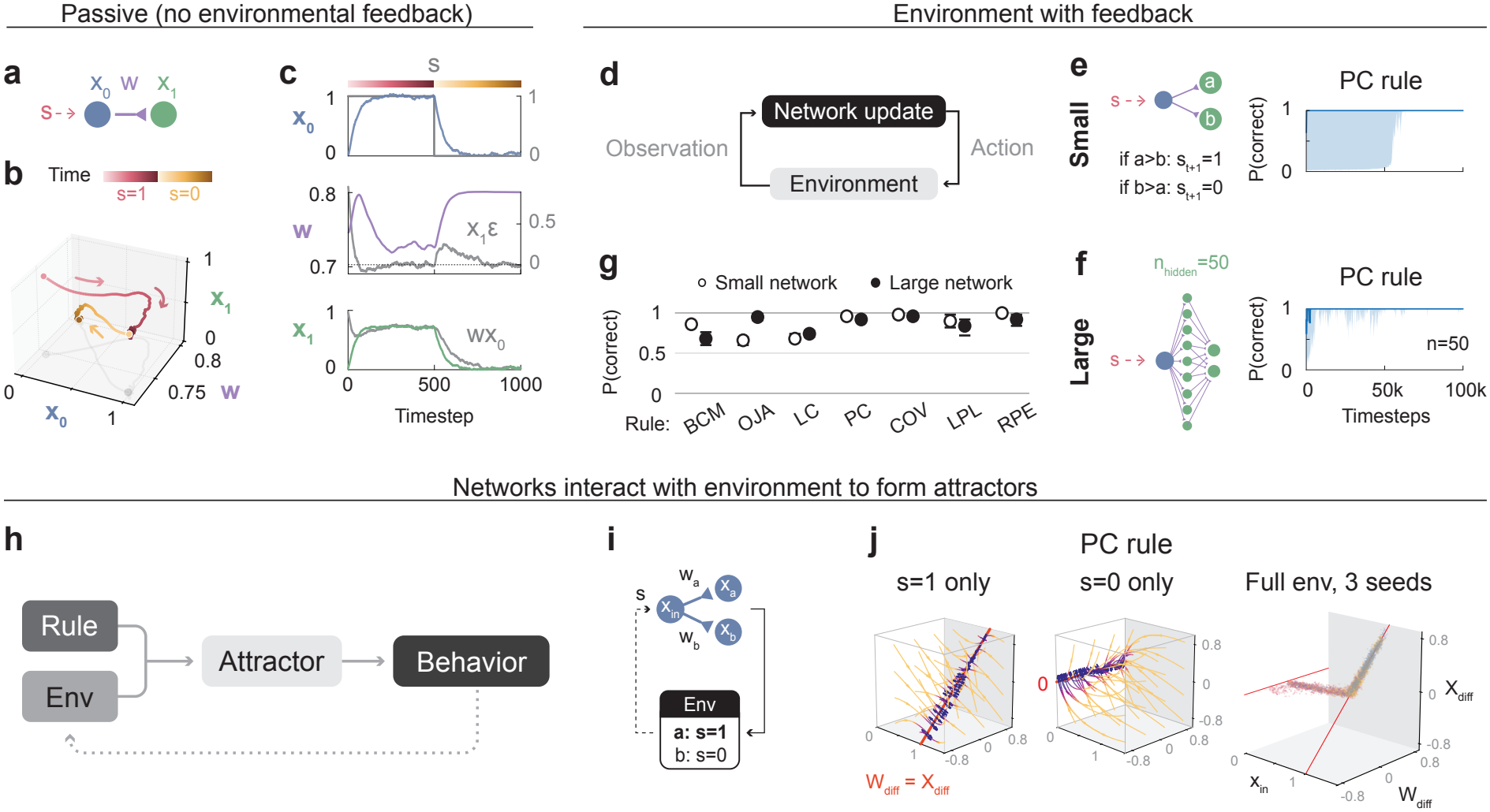
Local plasticity rules learn to seek signal in closed-loop environments. **a**, A two-neuron network with activities *x*_0_ and *x*_1_, a weight (*w*), and input signal *s*. **b**, Variables can be visualized as a single dynamical trajectory for a fixed input sequence, *s* = 1 for 500 timesteps and *s* = 0 for another 500 timesteps. Weight dynamics use the PC rule. **c**, Variables plotted individually over time for the trajectory in **b**. Also plotted are *x*_1_*ε* and *wx*_0_ because they determine how *w* and *x*_1_ update, respectively. **d**, Networks can be placed in environments that respond to actions at every timestep. **e**, Three-neuron networks learn to seek signal (*s* = 1) over time. A 100-step rolling mean is computed over each seed and median values are shown with 25th–75th percentiles. **f**, Same as **e** but for networks with 50 hidden neurons. **g**, Small (3-neuron) and large (53-neuron) networks learn to seek signal under all tested learning rules (0.5 is random). Points show the median over steps 80k–100k of P(correct) averaged across 50 seeds. Error bars, 95% bootstrap CI over seeds. In **e**–**g**, network hyperparameters are selected as the best-performing parameters on this metric, with *η* ≥ 10^−5^. **h**, Networks and their environments interact to form attractors, and these attractors determine behavior. **i**, The deterministic environment considered for panel **j**. Choosing *a* gives *s* = 1 and choosing *b* gives *s* = 0. **j**, PC attractors in network state space (*x*_in_, *W*_diff_, *X*_diff_) given constant input *s* = 1 (left), 0 input (center), and feedback in the responding environment of **h** (right). Red lines, lines of fixed points. Three seeds show overlapping occupancy of the attractor, which is the union of the two branches. *W*_diff_ = *w*_*a*_ *− w*_*b*_ and *X*_diff_ = *x*_*a*_ *− x*_*b*_ are the differences between the input weights and the activities of the two output neurons.

Sensory stimuli in passive environments (Fig. S1a,b) and rewards in interactive environments are both given as inputs (Fig. 1d–f). Passive environments test for the ability to learn unsupervised transformations of input patterns, while interactive environments test the ability to learn adaptive behaviors, often by seeking out reward in some sense. The difference between the two environment types (Table 1) is that only interactive environments respond to network actions through winner-take-all choices. That is, every timestep an action is chosen by an argmax on the network’s output layer, corresponding to the index of the neuron that is most active, and that index is mapped by the environment to a response.

### 2.2 All seven rules can seek signal, in parameter regimes that differ by rule and task

We evaluated every rule across previously developed representation learning and adaptive behavior environments (Methods). Rules OJA, PC and RPE involve three shared hyperparameters (activity update rate *α*, weight learning rate *β* and noise scale *η*) while rules BCM, LC, COV and LPL have an additional time constant *τ* (Methods). We swept several orders of magnitude of each hyperparameter in the small and large architectures from Fig. 1e-f, finding that learning rules could support performance on tasks for which they were not originally designed (Fig. S1– S3). For instance, RPE-networks learned feature selectivity while BCM-networks learned the two-armed bandit (Fig. S1a,c). The specific tasks a network could perform depended on its parameters (*α, β, η, τ*) as much as on its rule, and best-performing settings differed between rules as well as between tasks (Fig. S2–S3). We consider the implications of this in the Discussion.

In two-choice deterministic environments, all rules showed basic signal-seeking behavior (Fig. 1e–g). To see why, note that in every rule each weight update is multiplied by presynaptic activity (Eq. 1). When the chosen action produces input, it keeps driving the output that won, and the weights onto that output are updated so the same action tends to be chosen again. When the chosen action produces no input, the input neuron falls silent, and plasticity stops. The outputs, which are driven only through the input, relax toward zero, so the next choice is set by noise. Actions that produce input are therefore self-sustaining, while other actions revert to noise. This parallels the result of Loewenstein and Seung that plasticity driven by the covariance between reward and neural activity produces reward seeking and matching. ^17^ However, because updates here are gated by presynaptic input, the input itself plays the role of reward, and no reward term is needed. In comparison, the three-factor RPE rule reaches the same outcome through its explicit reward term.

To understand how local mechanisms produce behavior, it is useful to consider the dynamical system formed by the three-neuron PC network (Fig. 1h,i). Actions depend on the relative activity of output neurons, so only their differences matter for behavior. The coordinates we plot are thus *x*_in_, *X*_diff_ = *x*_*a*_ − *x*_*b*_ and *W*_diff_ = *w*_*a*_ − *w*_*b*_. The deterministic environment has two cases, *s* = 1 and *s* = 0. Setting both activity and weight update equations to zero in each case reveals two line attractors, ^47,48^ shown by the vector fields in Fig. 1j: for *s* = 1 the fixed points form the line *X*_diff_ = *W*_diff_ (with *x*_in_ = 1), and for *s* = 0 every point with *x*_in_ = *X*_diff_ = 0 is fixed. In the responsive environment, the combination of the two lines forms a bridge (Fig. 1j, right). A network’s learning rule thus interacts with an environment to form an attractor, which then determines downstream behavior (Fig. 1h), and we call these “world-in-the-loop” attractors. In the sections below, we use these attractor geometries to explain each behavior.

While all learning rules tested learn to seek signal, only some capture behavioral phenomena recorded in animals. Here we present a set of tasks that distinguish between rules. Near the parameters where each rule seeks signal, prospective configuration (PC) was the only rule to capture all phenomena tested. Other rules reproduce some of these phenomena elsewhere in their parameter spaces (Fig. S2–S4), so the comparison describes where each rule works well rather than every possible setting.

### 2.3 Reward magnitude scales the rate of learning

We first asked what determines learning rates in these networks. It has long been known that more intense stimuli drive faster learning; in classical conditioning, more intense conditioned stimuli produce faster acquisition, ^49^ although there the stimulus predicts an outcome rather than resulting from the animal’s actions. In classic models of associative learning, reward magnitude can influence the performance to which learning converges (the asymptote). ^50^ But Gong et al. ^32^ recently showed that across various tasks, mice learn faster with larger rewards, while the asymptote does not necessarily change (Fig. 2a,b).

**Figure 2:**
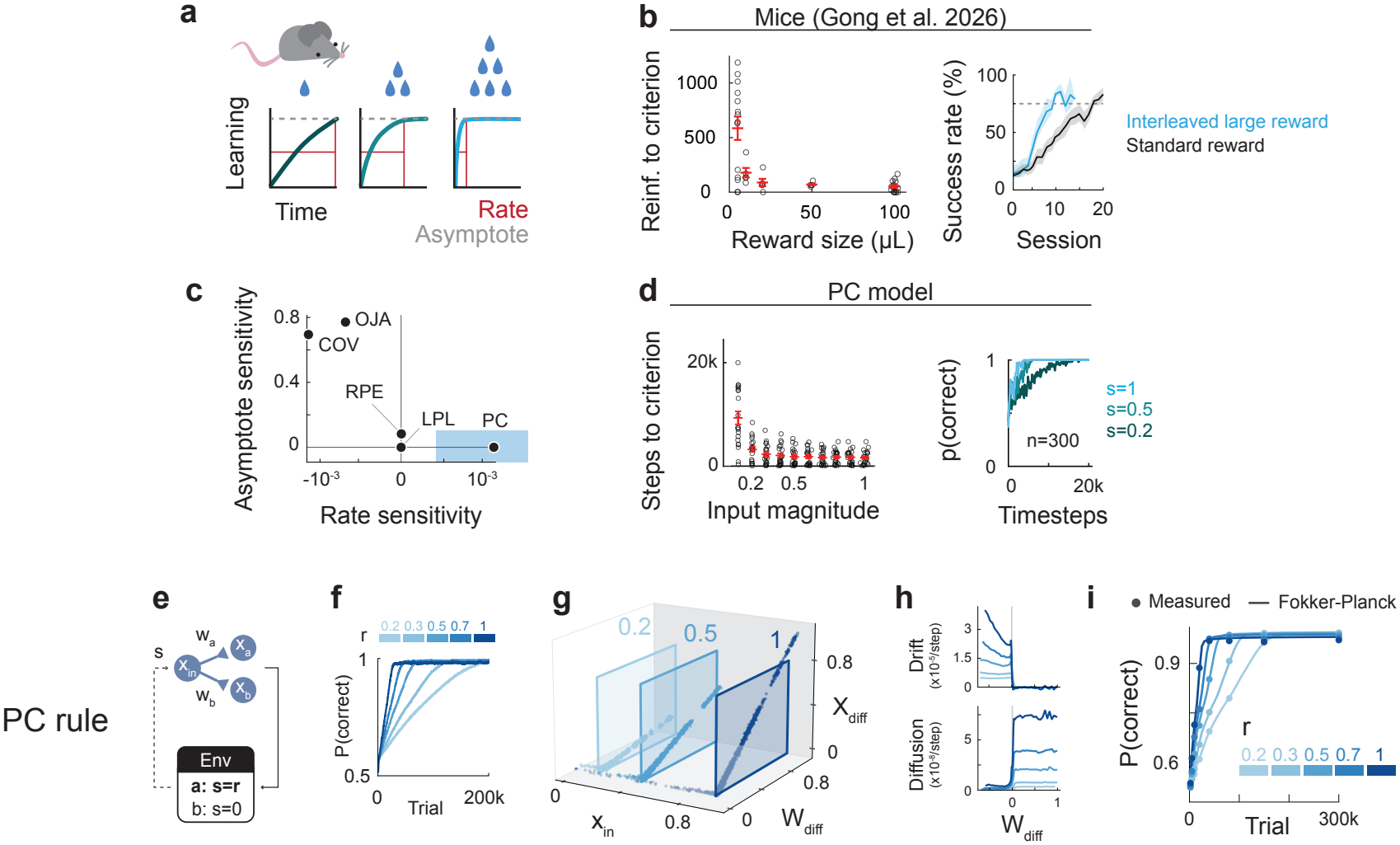
Reward magnitude scales the rate of learning. **a**, Cartoon: reward sizes influence learning rates but not asymptotes. Based on Gong et al. 2026. ^32^ **b**, Adapted from Gong et al. ^32^ Left: our reanalysis of hidden-target navigation data: reinforcements until each mouse’s smoothed target latency is 90% of the way from its initial to its trained level (Methods). Right: success rate across sessions in an effort-based joystick task, digitized from Gong et al. (2026), their Fig. 4B. **c**, Asymptote and rate sensitivities of all rules using best-performing hyperparameters in the deterministic environment (Fig. 1; see Fig. S4 for metric definitions). The shaded blue box indicates the region consistent with animal behavior, where learning rate is sensitive to input magnitude but the performance asymptote is not (Supplementary Note of Gong et al. ^32^). Note that many rules span asymptote and rate sensitivities for different hyperparameter combinations (Fig. S4). BCM and LC are not shown: in the large network they reach a signal-seeking plateau only at the largest input magnitude, and at smaller magnitudes their choices move toward the unrewarded arm, so no learning rate can be measured (Section S1). **d**, Sample efficiency of a PC model (large network) with different input magnitudes. Left: timesteps to criterion, the first 100-step bin at which median P(correct) across 20 seeds reaches 0.9. Circles are 20 independent batches of 20 seeds; red bars, mean *±* s.e.m. across batches. Right: median learning curves as in Fig. 1e (*n* = 300). *In PC, reward magnitude affects learning rate by modulating drift and diffusion along attractors*. **e**, The deterministic environment with input magnitude *r* for the rewarded action. **f**, Mean performance across 800 seeds (10 batches of 80) for different reward magnitudes. Larger rewards learn faster but reach the same asymptote. **g**, Attractor branches after learning for the simulations in **f**, in *W*_diff_–*X*_diff_ space (*W*_diff_ = *w*_*a*_ *− w*_*b*_, *X*_diff_ = *x*_*a*_ *− x*_*b*_). The slope approximately equals the reward magnitude *r*. **h**, Drift and diffusion along the attractors. Binned drift and diffusion of the weight-difference coordinate, per reward magnitude. Drift is positive while *W*_diff_ favors the unrewarded arm, is proportional to *r*, and vanishes once the network reliably chooses the rewarded arm. Both drift and diffusion grow with reward. **i**, A one-dimensional diffusion reproduces the learning curves. Circles: measured P(correct) at several timepoints (held-out seeds). Lines: the same quantity predicted without fitting to these curves, using the drift and diffusion estimated in **h** from separate seeds and integrating the Fokker–Planck equation forward from each reward level’s empirical initial *W*_diff_ distribution. Color, reward magnitude. The reward dependence of learning in **f**, including the common asymptote, is reproduced from a single scalar drive acting through the attractor’s geometry.

We placed networks with 50 hidden neurons in deterministic two-alternative forced choice (2AFC) tasks with varying input magnitudes. We computed the asymptote and rate sensitivities of each rule at its best-performing parameters from Fig. 1g and found that the rules spanned a space of sensitivities (Fig. 2c). At these parameters only PC showed rate but not asymptote sensitivity (Fig. 2c,d), matching animal data. This outcome is parameter-dependent, and examples of various combinations of rate and asymptote sensitivity can be found in all rules (Fig. S4), although not at the best signal-seeking parameters for rules other than PC.

In PC, this reward dependence follows from the geometry and dynamics along the attractor (Fig. 2e-i). For weights that favor the rewarded action, the attractor’s slope in *X*_diff_-*W*_diff_ space equals the reward magnitude *r* (Fig. 2g), so the same change in weight produces a larger change in output (and thus faster learning) at larger rewards. Learning is slow movement along the line attractor and can be described as one-dimensional drift and diffusion of *W*_diff_ (Fig. 2h). Because drift vanishes once a network reliably chooses the rewarded action, learning stops at the same performance for every reward magnitude.

With drift and diffusion estimated from separate networks, this description predicts the learning curves of held-out networks without fitting (Fig. 2h,i). A companion paper derives these drift–diffusion dynamics analytically for the three-neuron network and shows in simulation that the same line forms in the network with 50 hidden units used in Fig. 2.^48^

### 2.4 Choice allocation: probability matching and costly stochasticity

#### Probability matching

Once networks have learned, how do they distribute their choices? Animals are often less exploitative than is optimal, yet their randomness is sensitive to the environment. One consistent behavioral signature across species is known as probability matching, where in a 2AFC task, animals match the ratios of their choices to the ratios of reward probabilities ^33,34^ (Fig. 3a-c, showing mouse data from Bari et al. (2019) ^51^ and Beron et al. (2022) ^35^). This is optimal when rewards are baited, but clearly suboptimal when they are not, and animals probability match in both. Reward-maximizing agents do not match in the unbaited setting, which is the standard two-armed bandit, and instead converge on the better arm. ^9,52^

**Figure 3:**
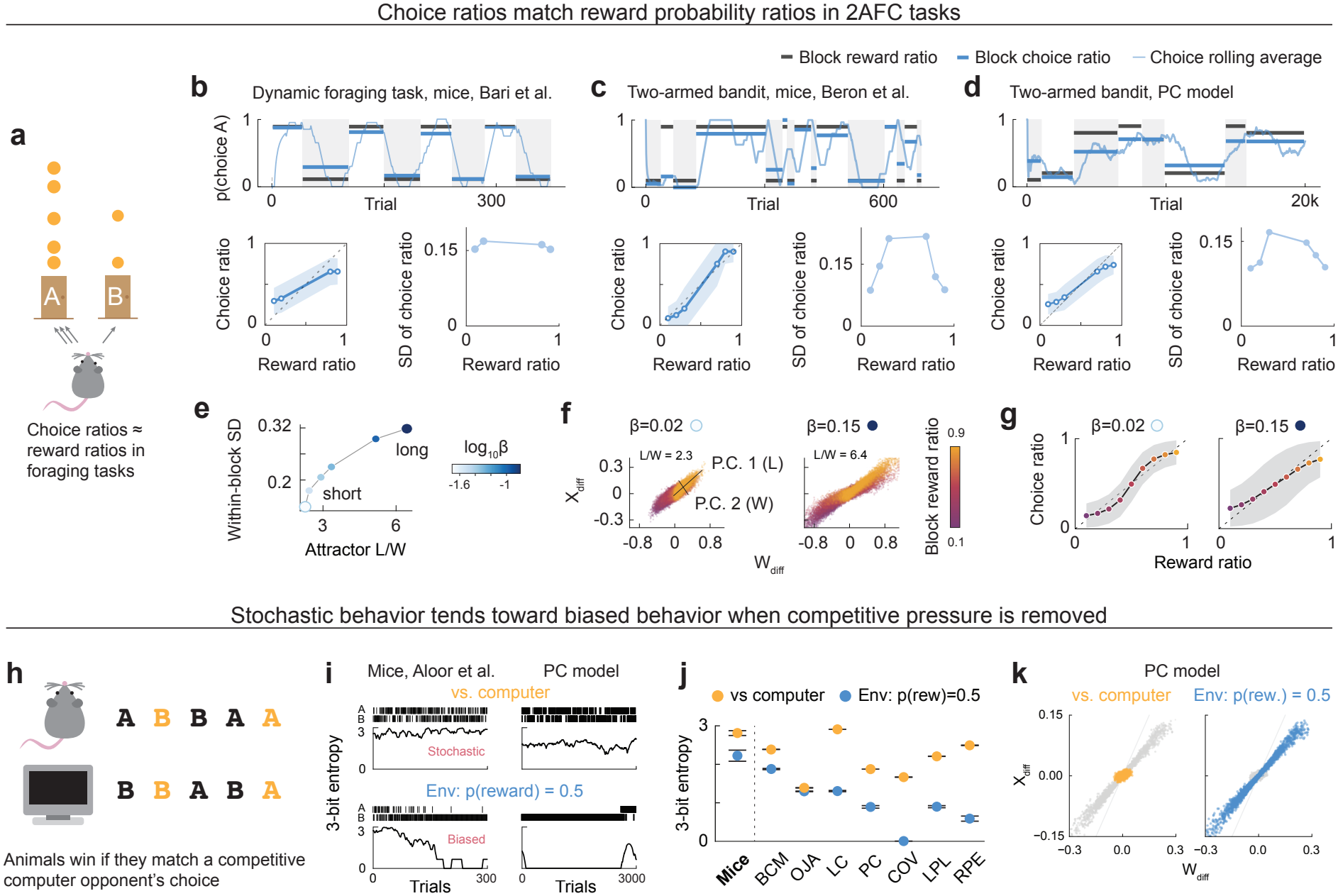
Plasticity rules probability-match, attractor length sets choice variability, and networks become more random under competition. **a**, Animals choose in proportion to reward ratios (probability matching). This is close to optimal in dynamic foraging (**b**) but not in the two-armed bandit (**c**,**d**), where always choosing the better side is optimal. **b**, Dynamic foraging (Bari et al. ^51^). Top: example session, rolling P(choice A) (20-trial window); black and blue bars, each block’s reward and choice ratios. Bottom: choice ratio vs. reward ratio (mean *±* s.d. across blocks, first 10 trials dropped; dashed, perfect matching) and s.d. of the block choice ratio. **c**, Two-armed bandit (Beron et al. ^35^), as in **b** (30-trial window). **d**, PC in the two-armed bandit, as in **b** (1,000-trial window; first 2,500 trials of each block dropped; mean block length 5,000 trials). Parameters were fit to **c**, but other parameters and rules also match (Fig. S1–S3, Fig. S5). *In PC, attractor length sets choice variability*. **e**, Within-block choice s.d. (reward ratio 0.5) vs. attractor length-to-width ratio *L*/*W* across a *β* sweep. **f**, Attractors of a short (*β* = 0.02) and a long (*β* = 0.15) agent in *W*_diff_–*X*_diff_ space (*W*_diff_ = *w*_*a*_ *− w*_*b*_, *X*_diff_ = *x*_*a*_ *− x*_*b*_), colored by block reward ratio. *L* and *W*, extents along the first two principal components. **g**, Both agents match on average but differ in variability (shading, within-block s.d.; 0.20 vs. 0.32 at reward ratio 0.5). Baited task, 40 seeds. *Stochastic behavior becomes biased when competitive pressure is removed*. **h**, Matching pennies: the agent is rewarded when a computer opponent fails to predict its choice from its history. **i**, Example choices and 3-bit entropy. Mice (Aloor et al. ^36^): 300 trials, 30-trial window; median session against the computer, a clearly biased session where the environment is defined as p(reward) = 0.5. PC: 3,000 trials and a 300-trial window, scaled by 10 for its slower timescale; median seed (Methods). **j**, Entropy per rule and for mice (mean *±* s.e.m.; models, 50 seeds, settled phases; mice, *n* = 7, each mouse’s mean over sessions of each type). Entropy is higher against the computer for every rule (*p <* 10^−13^, one-sided paired Wilcoxon, Holm-corrected) and for mice (*p* = 0.008). **k**, PC attractors over the last 5,000 steps of a late phase (50 seeds; grey, the other condition). Attractors are roughly point-like against the computer and lengthened without competition.

However, matching can be reproduced by models fit to behavior, including reinforcement-learning models with a stochastic (softmax) choice rule ^53^ and local estimates of recent reward income, ^34^ and has been shown to arise from some forms of reward-dependent plasticity.^17,54^ To see whether local networks generally capture this behavior, we tested all rules in both baited and unbaited environments. In the small network, four rules (RPE, PC, COV and LPL) probability-matched in both environments and BCM in the baited environment only. In the large network, RPE, PC, COV, LPL and OJA matched in both (Fig. 3d; Fig. S1–S3 and Fig. S5; Methods).

Mechanistically, we find that attractor geometry correlates with choice variability in the PC networks (Fig. 3e). The weight learning rate *β* modulates the length-to-width ratio of attractors (Fig. 3f), and these attractors mediate behaviors that probability-match equally well on average but have different amounts of within-block variability (Fig. 3g). Within the tested parameter range, faster weight learning therefore leads to longer attractor geometries, which generate more variable choices (Fig. 3e).

#### Costly stochasticity

Animals compete for resources, ^55^ and in competitive interactions being predictable can be costly. Predictable prey are easier for predators to catch, and predictable predators are easier for prey to escape. In the laboratory, such interactions are often studied with games. ^56^ One game known as matching pennies ^57^ asks each player to pick one of two options on every turn; one player is rewarded when choices match, and the other when they do not. When both players are rational, the optimal strategy is to choose at random. Mice playing a computer opponent that exploits patterns in their choices learn to be highly stochastic (Fig. 3h; Methods), but when the opponent is replaced by a coin flip, they quickly become biased, suggesting that acting stochastically carries a cost ^36^ (Fig. 3i).

We alternated competitive and non-competitive environments to see how networks would behave, choosing for each rule the parameters that gave the largest change in choice entropy between the two once behavior had settled (Methods). Every rule chose more randomly against the computer opponent than when competitive pressure was removed (Fig. 3i,j), but by very different amounts: OJA barely changed (0.09 bits), whereas COV went from 1.66 bits to choosing one side exclusively. Most networks were less random than the mice against the computer: mice reached close to the maximum of 3 bits (2.81 ± 0.06, mean ± s.e.m. across 7 mice), as did LC (2.90), but PC reached 1.87 due to its choice following a weight that drifts slowly relative to the timescale of individual choices. Attractor dynamics in the PC network revealed that while biased behavior was mediated by line-like attractors as in Fig. 1–3, high-entropy random behavior came from compressed, point-like attractors instead (Fig. 3k).

### 2.5 Temporal structure: discrete states and heavy-tailed run lengths

#### Discrete behavioral states

Beyond the overall distribution of choices, animal behavior has temporal structure. Hallmarks of animal behavior include persistent choices and autonomous switching between distinct behavioral states. ^37,38^ Using the environment from Fig. 1d–f, we looked for evidence of discrete explore and exploit states by fitting two-state hidden Markov models (HMMs) and comparing the fits to no-switching models (Fig. 4a; Fig. S6 for details). We found that five of the seven rules, LC, PC, COV, LPL and RPE, showed discrete bistability (Fig. 4b), and in PC, this bistability is mediated by the two attractor branches in Fig. 1j.^47^ Networks do not settle on the preferred action, as exploit episodes are roughly uniformly distributed over time even on long time horizons (Fig. 4c; 5 million steps in Fig. S6). The intriguing conclusion is that several simple learning rules, even in toy models and deterministic environments, allow networks to balance autonomously between discrete states of exploration and exploitation.

**Figure 4:**
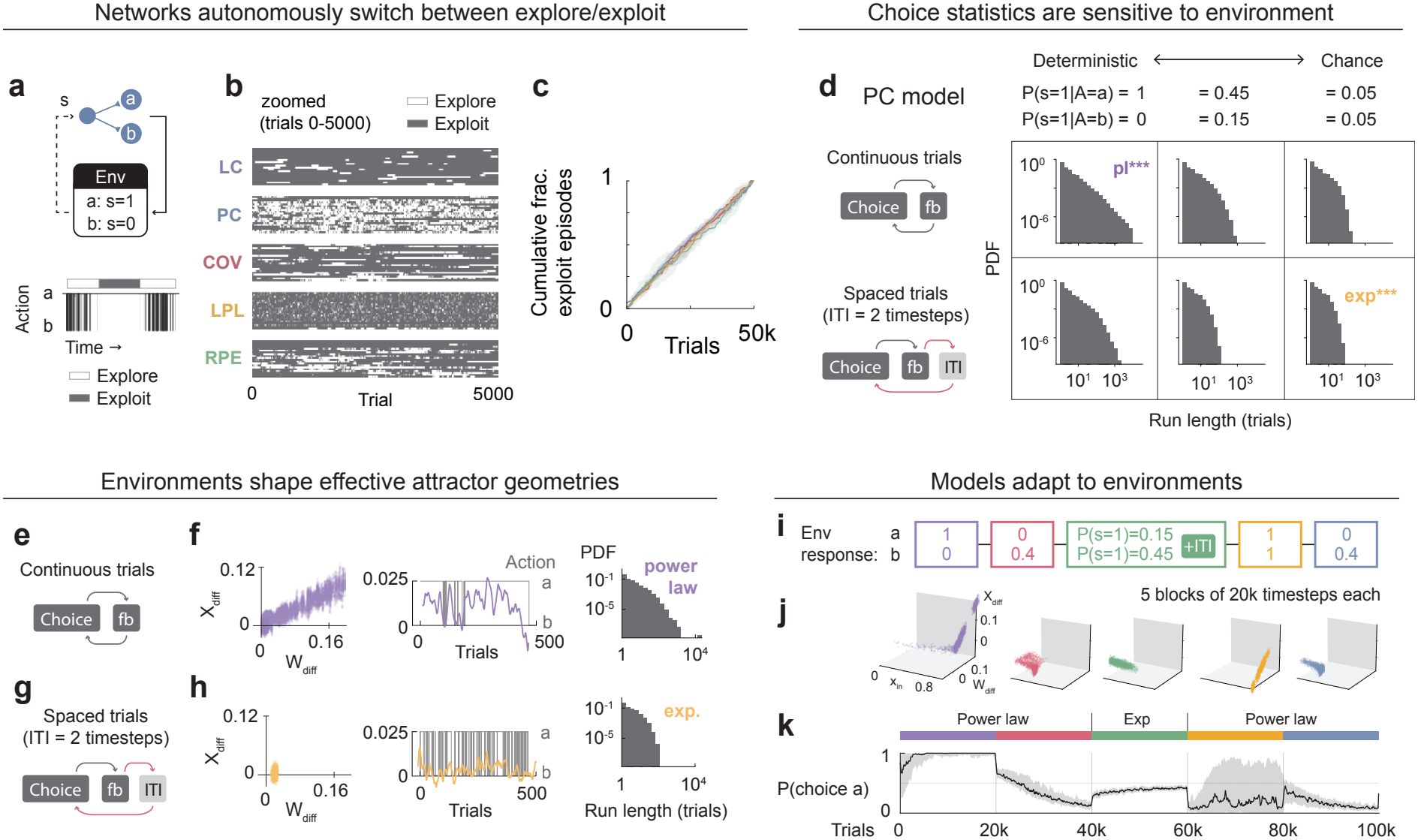
Discrete explore–exploit states, environment-dependent run lengths and adaptive attractors. **a**, Cartoon of the deterministic environment and visualization of explore/exploit states. **b**, Discrete explore/exploit modes recur at steady state in most rules. Each panel uses a representative discretely bistable parameter set per rule, if available (Fig. S6), selecting the parameter set whose fitted two-state HMM switched most consistently across 20 simulated seeds (*T* = 50k trials). Trial states are recovered by Viterbi decoding (shown for trials 0–5,000 in 20-trial bins), with one row per seed, sorted top to bottom by total exploitation time. **c**, Cumulative count of exploit-state episodes started across all 50k trials, normalized by each seed’s final total to allow direct comparison across rules (Fig. S6). Bold line, median across seeds; shaded band, interquartile range. All five valid rules climb linearly rather than tapering, indicating that models do not settle into static action equilibria. **d**, Environmental conditions alone can affect the shape of run-length distributions. Examples of transitions between run-length distributions that depend on the continuity of environmental feedback or the probability of action-based feedback. Baited two-armed bandit, 120,000 trials. Vuong likelihood-ratio test (power law vs. exponential ^58^); *p <* 0.05 for a call, n.s. otherwise; ^***^*p <* 0.001. Hyperparameters from Fig. 5. *In PC, environments shape effective attractor geometries*. **e**, Continuous trials (ITI = 0): every timestep is a choice followed by feedback (fb). **f**, Continuous environment. Left: attractor cloud in *W*_diff_–*X*_diff_ space (*W*_diff_ = *w*_*a*_ *− w*_*b*_, *X*_diff_ = *x*_*a*_ *− x*_*b*_) over a long (360k-step) run, which spends 76% of trials on the good arm and elongates far along *W*_diff_ (line-attractor-like). Middle: *X*_diff_ and the corresponding action over a representative window. Right: pooled run-length distribution (PDF, log–log); power law, *p <* 0.001 (32,591 pooled runs, longest 22,900 trials). **g**, Spaced trials, with an inter-trial interval of two timesteps (ITI = 2). **h**, As in **f**, for the spaced environment. The run (64% good arm) stays confined to a narrow band near the origin (point-attractor-like), and run lengths are exponential, *p <* 0.001 (75,194 pooled runs, longest 80 trials). In **e**–**h**, networks use the settings from **d** but with lower activity noise (*η* = 0.002); the environment is a baited bandit, *P* (*s* = 1 | *A* = *a*) = 0.45, *P* (*s* = 1 | *A* = *b*) = 0.15; Vuong test as in **d**. *Models adapt to environments*. **i**, Networks are placed in an environment that changes every 20k timesteps, with deterministic (blocks 1, 2 and 5), probabilistic (block 3, with an ITI) and static responses (block 4). **j**, Networks form attractors suited to the current environment, shown in (*x*_in_, *W*_diff_, *X*_diff_) for each block. **k**, Choice probabilities for 48 seeds (median with 25th–75th percentiles) show that networks respond appropriately to each new environment block. The bar above marks the run-length distribution in each block (power law or exponential).

#### Heavy-tailed run-length distributions that are sensitive to environments

Many animals, from nematodes to humans, move with heavy-tailed run-length distributions known as Lévy flights. ^41,59^ Proposed explanations range from adaptation ^42,60^ and sensory perception ^61^ to optimal search ^62^ (although whether Lévy flights are optimal remains debated ^63^).

We find that simple synaptic mechanisms are sufficient to robustly capture two features of foraging: heavy-tailed run lengths and their dependence on the environment. Here run-lengths are defined as the number of consecutive choices of the same action. Using the PC model as an example, we find that heavy-tailed run-length distributions emerge naturally in the deterministic environment of Fig. 1d–f and Fig. 4a–c (Fig. S2–S3). Networks transition between heavy- and light-tailed run-length distributions depending on the statistics of the environment (Fig. 4d), a trait also seen in animal foraging data. ^59^ Here networks show heavy-tailed distributions when feedback is consistent and deterministic, and exponential distributions when feedback is temporally spaced and random.

In PC, environments change run-length distributions by modulating attractor geometries. When trials are continuous, line-like attractors form in the network state space (Fig. 4e,f), and random walks up and down the attractor cross the decision boundary with power-law distributions (Fig. 4f). This is consistent with the heavy-tailed first-passage times of random walks. ^64^ In contrast, when trials include inter-trial intervals (Fig. 4g), attractors contract until they look more like noise about a point (Fig. 4h). In these cases, boundary crossings have exponentially distributed timings (Fig. 4h).

#### Adapting to changing environments

We also tested the adaptability of these attractors by placing networks in continually changing environments (Fig. 4i–k). Attractor geometries (Fig. 4j) and behaviors (Fig. 4k) readily change with the environment, reproducing several of the behavioral phenomena described above, including transitions between heavy- and light-tailed run-length distributions, in an online, continuous, closed-loop setting.

### 2.6 Memory when the environment changes

#### Interference-dependent forgetting

Last, we asked what happens to what networks have learned when the environment changes. McGeoch ^44^ argued that forgetting did not come from the passage of time but from events that interfered with stored memories, and to see whether local learning rules captured this now well-known phenomenon, ^43^ we placed trained networks in three different conditions: no input, constant input, and reversal (Fig. 5a,b). PC, COV, LPL and RPE stored the trained preference in their weights, and retained more of it with no input than with constant input (Fig. S7 and Table S1), showing that local rules can exhibit interference-dependent forgetting. BCM and OJA differed between conditions, but most of their networks stored almost no weight difference during training (median *W*_diff_ ≈ 10^−4^), so this difference reflects constant input driving *W*_diff_ negative rather than retention. LC did not differ.

**Figure 5:**
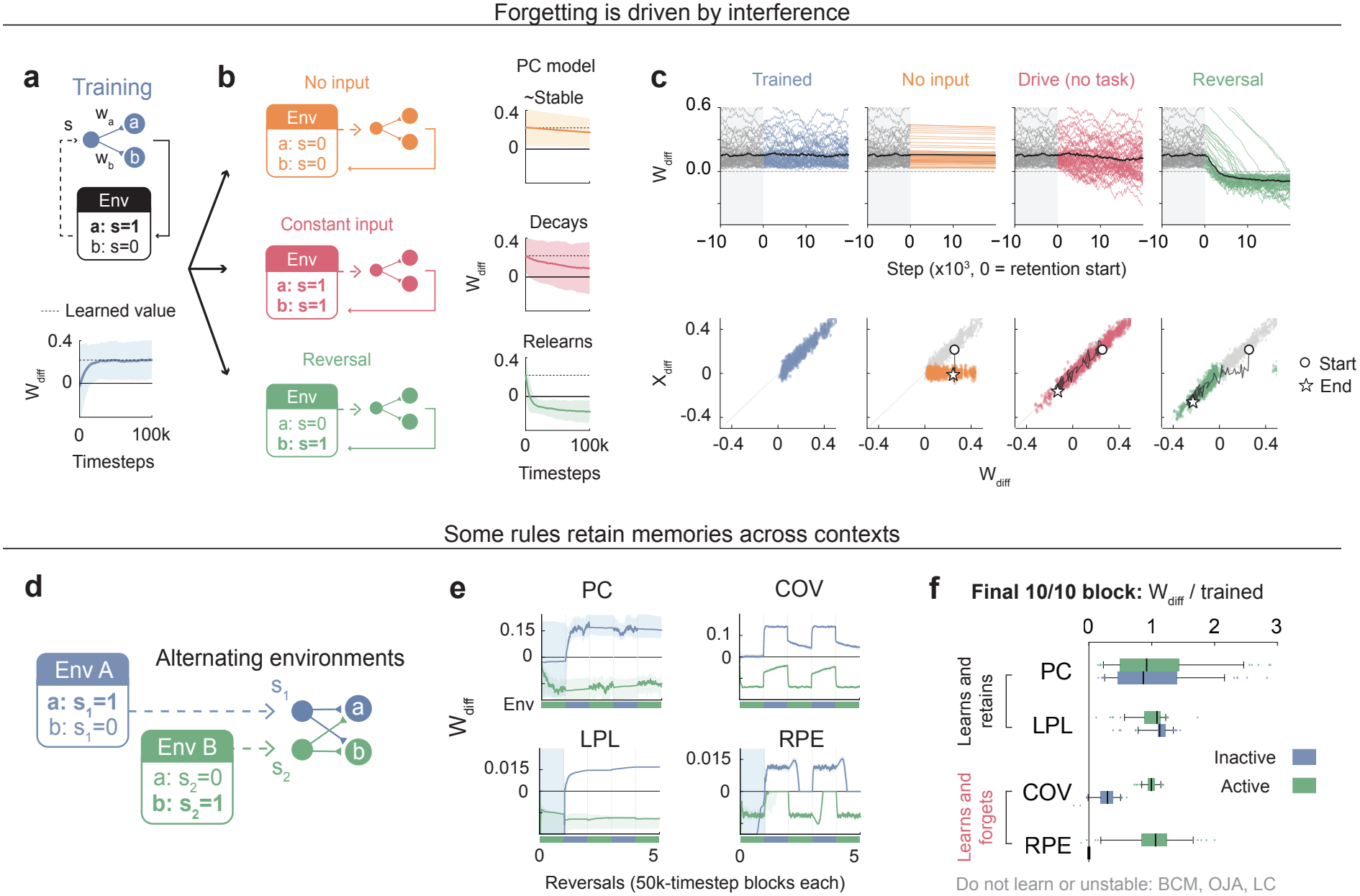
Rules show interference-dependent forgetting and retain memories across contexts. **a**, In each of three two-stage tasks, networks are first exposed to a deterministic environment with one rewarded choice. We use PC as an example, with the best-performing parameters on the deterministic task (Fig. 1g), then filtered for performance on the forgetting task (see Methods, and Fig. S7 for other rules). Bottom: networks learn over 100k timesteps (*n* = 250). We plot the mean difference between weights, which shows the preferred action. Shaded regions, s.d. **b**, Top: in one task, network dynamics continue to evolve while receiving 0 input. Middle: in a second task, networks receive constant input without task structure. Bottom: in a third task, networks must respond dynamically to the environment by learning a reversal. **c**, Attractor transitions underlie interference-driven forgetting in PC. Top: *W*_diff_ = *w*_*a*_ *− w*_*b*_ (*a*, the originally rewarded arm) for 50 individual seeds (thin lines) and the median (black) over the last 10k training steps (grey) and 20k steps of each condition: continued training (trained), no input, constant input without a task (drive), and reversal. Bottom: the same runs in *W*_diff_–*X*_diff_ space (*X*_diff_ = *x*_*a*_ *− x*_*b*_). Grey, trained attractor cloud; colour, cloud at the end of each condition; line, one example seed from start (circle) to end (star). Retained memories (no input) are kept by rotating the attractor into the weight dimension. Constant input allows diffusion across the entire attractor branch, and reversal moves the network onto the opposite branch. **d**, Next we test the case where inputs for different contexts are received by different neurons. **e**, Weight differences for each context across the first 5 of 10 reversal blocks (*n* = 100). Lines are medians and shading is the 35th–65th percentile range. PC and LPL show little to no forgetting between active contexts. COV and RPE show forgetting, with RPE almost fully forgetting between blocks. However, RPE maintains good performance in the reversal task (Fig. S8). **f**, Relative weight of the active (green) and inactive (blue) context at the end of the last of 10 blocks, as a fraction of each rule’s achievable trained weight. Boxes, IQR; centre line, median; whiskers, 5th–95th percentiles. A rule retains memory if *≥* 95% of simulations have an Env A preference *>* 0: PC (100/100) and LPL (99/99) do (one-sided binomial test, Holm-corrected *p* = 0.024 for both); COV (94/100) and RPE (18/100) do not. Only rules where a majority of trials had a trained scale between 0.005 and 1.0 (above which weights diverged) are included.

In PC, the interference-driven forgetting in Fig. 5 can be visualized and understood as four distinct attractors in activity–weight space. A trained attractor is a branch that lies only on the responsive side of *W*_diff_ (Fig. 5c, trained). When input is 0, the attractor rotates into weight space, preserving the memory while activities throughout the network are 0 (Fig. 5c, no input). If constant input or a reversal task is presented instead, the network drifts along the line attractor into an unlearned or reversal-learned state, respectively (Fig. 5c, drive and reversal).

#### Memory retention across contexts

We next asked whether memories survive when networks alternate between contexts. We had networks experience alternating blocks of environments that fed into different context (input) neurons (Fig. 5d). This idealizes context-dependent routing as suggested from neural data ^65–67^ and by prior models. ^68^

Despite being able to relearn tasks each block, the three-factor RPE rule retained almost none of the inactive context (18% of simulations above zero), unlike the local PC, COV and LPL rules (100%, 94% and 100%; Fig. 5e,f; Fig. S8). COV also decays between the 50k-timestep blocks (Fig. 5e,f) because unlike other rules, it requires an explicit weight decay term for stability.^17,69^ PC and LPL, on the other hand, were remarkably stable.

## 3 Discussion

We have shown that local Hebbian-like learning rules are sufficient for a wide range of adaptive behaviors. In PC, these behaviors arise from attractors in activity–weight space that are not specified in advance but form from the interaction of network and environment. Given the long history of local learning rules, why was their capacity for adaptive behavior somewhat over-looked? Several papers have shown Hebbian-like rules performing goal-directed behavior, ^18–20^ but to our knowledge, this is the first systematic comparison showing that canonical local rules, as a group, can learn to seek signal in closed-loop environments.

We see two reasons for this gap. First, reward learning was usually studied in environments where rewards arrive long after the actions that caused them. This goes back at least to Marvin Minsky’s SNARC (1952) ^70,71^, which simulated rats running a maze, and these harder environments did require global signals and eligibility traces. ^9,14^ But many environments respond quickly, both in nature and in the lab—two-alternative forced choice tasks, for instance, are a staple of behavioral neuroscience, and reward animals quickly after decisions. A second reason is that reward was often assumed to be a special global signal, which local rules didn’t have. But the rewards that animals receive frequently have sensory components, and studies have shown that sensory components alone can drive behavior. ^23,24^ In a real brain, diffuse modulators may be triggered by these sensory signals, but sensations can also enter networks as ordinary inputs, as we model here. ^21,22^

### 3.1 Reward-seeking without direct optimization

Current frameworks for reward-seeking behavior emphasize explicit optimization. They also emphasize the need for neuromodulators that are distinct from sensory input. ^9,13^ Our results suggest that these choices are not always necessary, and instead, one role of broad neuromodulation may be to shape and direct local learning, which is capable of signal-seeking behavior on its own. ^2,72^

Is the input neuron simply a reward signal under another name? We believe this framework differs from reward-based ones for three reasons. First, the gate on plasticity is the presynaptic activity of each synapse, a factor already present in every two-factor rule (Eq. 1), so no additional modulatory factor is needed. Second, the gate is local. In the large network, synapses onto the output layer are gated by hidden-layer activity, which reflects reward only through propagated activity. In contrast, a three-factor signal is broadcast identically to all synapses. Third, the two approaches lead to different behavior, which we can see from comparing the RPE rule to local rules. Unlike PC and LPL, the RPE rule nearly erases the inactive context when contexts alternate (Fig. 5e,f), consistent with its error signal reaching every synapse rather than only those of the active context. We also note that under our framework, preferred inputs can still be distinguished from nonpreferred ones by circuit properties such as connectivity strengths or individual neural learning rates. ^47^

The models shown here are not designed to optimize for reward, as reinforcement learning models do; nor are they fit to behavior, as many networks trained with backpropagation are ^73^ (see Brunton et al. ^74^ for a cautionary example of the latter). Like animals, our models learn online, with no separate training and test phases, and they naturally allow for “suboptimal” behaviors that animals show—as from a purely reward-seeking perspective, probability matching in two-armed bandits and continued exploration in static environments are suboptimal. Common explanations for such behaviors invoke cognitive biases or evolutionary pressures in dynamic worlds, ^75,76^ but as others have written, such explanations are often constructed post hoc, and with enough room in the assumed environment or objective, almost any behavior can be rationalized as optimal. ^77,78^ We take a different approach, asking which behaviors can emerge from which mechanisms and a handful of parameters, and instead of competing with evolutionary accounts, these mechanisms may be the way in which these evolutionary pressures act.

This framework also leads to a different notion of reward than Markov decision processes (MDPs), a standard treatment of agent-environment interactions. ^9^ In an MDP, reward is a scalar defined separately from the stimulus. Here, reward is the set of inputs an agent learns to seek out and is an emergent property of the agent–environment interaction, linking our work to cybernetics and parts of control theory that treat behavior as a way of regulating inputs and internal states. ^79–81^ While others have applied this perspective to neural systems in the past, ^18,82–84^ here it emerges from standard leaky integrator neurons and canonical Hebbian rules. As in Moore et al. (2024), ^84^ noise is essential in our framework (Li & Erlich ^48^), and importantly, noise does not just make behaviors noisier—increasing neuronal noise can even make behaviors more consistent (Fig. S2a). This further connects to work showing that noise can improve information transfer in biological systems. ^85,86^

### 3.2 Implications

Within a single rule, a few parameters (learning rates and noise) determine the tasks a network can perform. The same rule can support either representation learning or signal-seeking, which could explain the non-random variation in neural properties across brain regions. ^87^ It also may reflect the No Free Lunch theorems ^88^, in that if no single parameter setting is best for every task, a brain of differently tuned regions could be a good solution to natural intelligence while still using the same family of learning rules. Alternatively, neuromodulators may temporarily change parameters to steer networks toward specific goals. ^47,72,89^

Another question to examine is whether any given attractor in the brain is sustained by neural dynamics alone or by neural dynamics with synaptic plasticity.^90^ Line attractors are found in the oculomotor integrator, decision-making circuits and the hypothalamus, ^91–94^ and most models of these continuous attractors have static weights, either hand-tuned ^95^ or trained and frozen. ^92^ Models incorporating plasticity usually tune networks toward a specified attractor. ^96,97^ Hebbian rules have been shown to self-organize continuous attractors, but these earlier models have included training phases in non-responsive environments. ^98^ In contrast, our attractors have no target and change with online experience (Fig. 5c and Fig. 4j). It would be useful to probe which attractors in the brain are relatively static and which may emerge from environmental interactions. In either case, our model offers an explanation of how such attractors may be learned in the first place.

Our framework suggests experimental tests at the neural and behavioral levels. At the neural level, many behaviors should arise from learned, flexible attractors shaped by the network and its environment. These attractors live in activity–weight-difference space, which is hard to observe directly, but signatures should appear in neural activity along a decision axis. ^99^ Random-walk-like drift suggests a line attractor, while fluctuations about a fixed point suggest a point attractor. We have analyzed this geometry in detail only in the smallest network, where it can be visualized directly, but a companion paper shows that the same line attractor forms in a network with 50 hidden units. ^48^ Rules differ in their geometries as well; for instance, PC predicts stable line attractors while RPE and COV predict greater decay (Fig. 5). Such signatures could help distinguish between candidate learning rules in neural systems.

At the behavioral level, PC networks predict that shorter intervals between choices produce heavier-tailed run lengths. With continuous trials, biased random walks along line attractors produce power-law run lengths, while inter-trial intervals shrink the attractor and make run lengths exponential (Fig. 4e–h). This matches reported tempo effects on persistence, ^100^ and varying inter-trial intervals within a task would be a direct test of this prediction, as would looking for approximate line or point attractors that directly mediate foraging behaviors.

### 3.3 Limitations and future directions

The primary limitation of our model is that we study one rewarded input at a time; we do not compare multiple stimuli of different value within one network, and we do not model aversive behavior, which may require inhibitory neurons. Subsequent research can delve into more complex environments, observation spaces, and behaviors, along with the neural architectures that would make them possible. Similarly, more theory is needed on how architectures, parameters and environments shape attractors and behavior. Behaviors are qualitatively similar in the small and large networks here. The reduced coordinates (differences in weights and activities between output neurons) can be defined in larger PC networks, ^48^ but we have derived attractors only for the smallest network. Deriving them for other architectures, learning rules and environments is left to future work.

Otherwise, our models are deliberately minimal, making them tractable for theory and proof-of-concept simulations. They leave out many biological features, including additional timescales of dynamics, adaptation, neuromodulation, homeostasis, and interactions between multiple agents, each of which is a possible new direction. Models also choose actions at short intervals and are rewarded quickly, so harder environments, with delayed or sparse rewards, are a natural place to add neuromodulation (with local plasticity acting as an adaptive underlying substrate). And to handle environments with both complex sensory processing requirements and strongly motivating signals, networks could combine different parameters in different modules or include gradients of parameters, ^101^ to test whether a single autonomous system can solve tasks that require both representation learning and reward-seeking behaviors.

### 3.4 Conclusion

The models we have presented contain few components, none of which are particularly novel. They consist of leaky integrators, noise, winner-take-all dynamics and local synaptic updates. That such simple systems learn to seek signal at all is remarkable. That they also show signatures of animal behavior, some of which neuroscientists have long been interested in, suggests that a surprising amount of adaptive behavior may rest on the basic machinery of neural circuits and their coupling to the world.

## 4 Methods

### 4.1 Network model

#### Architecture

Networks are feedforward rate networks with an input layer, optional hidden layers and an output (“motor”) layer. We write *W*_*ij*_ for the weight from presynaptic neuron *j* to postsynaptic neuron *i*. A fixed binary mask T specifies connectivity: *T*_*ij*_ = 1 if *j* lies in some layer *l* and *i* in layer *l* + 1 (all-to-all between adjacent layers), and *T*_*ij*_ = 0 otherwise; only the masked weights 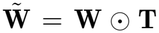 affect the dynamics. All neurons are excitatory, and input neurons receive no synapses. For the adaptive-behavior tasks we use two architectures: a *small* network, in which one input neuron projects directly to two output neurons (3 neurons, 2 plastic synapses), and a *large* network with one hidden layer of 50 neurons (53 neurons, 150 synapses). For the representation-learning tasks the input layer has one neuron per stimulus dimension (37 for feature selectivity, 2 for slow features) and there is a single output neuron. The memory retention task uses two input and two output neurons (Fig. 5d–f). Unless stated otherwise, activities and weights are initialized independently from a uniform distribution *U*(0, *κ*) with *κ* = 1, and running statistics (below) are initialized to *κ*.

#### Activity dynamics

Each neuron’s activity is predicted by its external input plus its presynaptic input, 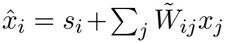, where *s* is non-zero only for input neurons. At every timestep activities take one gradient step of size *α* on the prediction loss 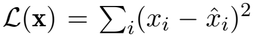, with 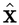 held fixed (a stop-gradient on the presynaptic term), x ←x −*α* clip(∇_x_ *L*, −10, 10). Written out, this is an Euler step of the leaky integrator in Eq. 2,

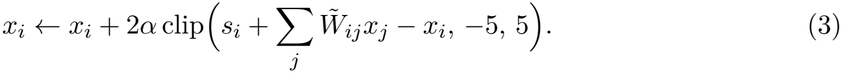

Independent Gaussian noise is then added and activities are rectified,

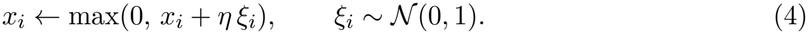

Input neurons receive no synapses and therefore relax toward *s*_*i*_. In some analyses activities were additionally capped at 3 after Eq. 4; this is stated per figure below and in Table S2.

#### Action selection and the closed loop

In two-choice environments the action at each timestep is the output neuron with the larger activity, *a*_*t*_ = arg max_*k*_ 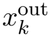, with ties broken uniformly at random. Each timestep proceeds as follows: (i) the action is read from the output activities at the end of the previous step; (ii) the environment returns a scalar input *s*_*t*_ (the “reward”), which is applied to the input neuron, s = (*s*_*t*_, 0, …, 0); (iii) activities are updated once (Eq. 3); (iv) noise and rectification are applied (Eq. 4); (v) weights are updated from the post-noise activities (Eq. 5); and (vi) running statistics are updated. Choices are made at every timestep, so “trial” and “timestep” are synonymous unless an inter-trial interval is present. Activity noise (and tie-breaking) is the only source of stochasticity in choice. In the representation-learning tasks the environment ignores the output and supplies a stimulus vector s instead.

### 4.2 Plasticity rules

All rules share the update

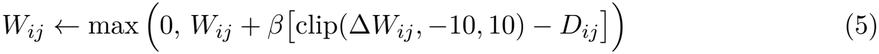

with weight learning rate *β* and a non-negativity floor. Each rule specifies only Δ*W*_*ij*_; the decay term *D*_*ij*_ is zero except for COV. Below, *x*_*j*_ is presynaptic and *x*_*i*_ postsynaptic, and running averages are exponential with rate *τ*.

#### Bienenstock–Cooper–Munro (BCM).^**27**^

Hebbian coactivity is compared with a sliding threshold set by the running mean activity *µ*_*i*_,

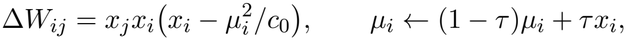

with *c*_0_ = 0.1 (the sliding-threshold normalization of Law & Cooper; the original uses *c*_0_ = 1).

#### Oja’s rule (OJA).^**28**^

A Hebbian term with multiplicative normalization,

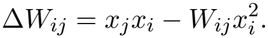

#### Law & Cooper (LC).^**29**^

The threshold *θ*_*i*_ is the running mean of the squared activity and also normalizes the update,

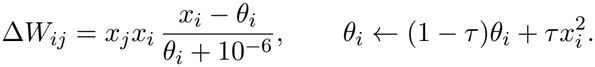

#### Prospective configuration (PC).^**30**^

Presynaptic activity multiplies the postsynaptic prediction error *e*_*i*_,

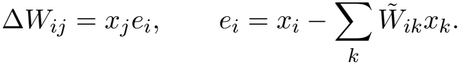

This is the negative gradient of the prediction loss *L* defined above with respect to W (up to a factor of 2) for every neuron that receives synapses, so activities and weights descend the same loss apart from a stop-gradient. Note that neuron activity updates differ here from Song et al. ^30^ in two ways: first, activities are not allowed to settle, but instead are coupled with the weights as in Li et al. ^47^. Second, activity updates do not minimize the exact prediction error derived from the weights. They differ by a stop-gradient (see *Activity dynamics*, above) that prevents postsynaptic activities from directly influencing presynaptic activities.

#### Covariance rule (COV).^**17**,**69**^

Postsynaptic activity multiplies the deviation of presynaptic activity from its running mean, with a soft upper bound on the weights,

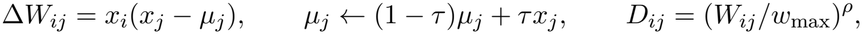

with *w*_max_ = 1 and *ρ* = 5. We use the unmodulated (two-factor) form throughout.

#### Latent predictive learning (LPL).^**31**^

A variance-normalized Hebbian term is combined with a predictive term that penalizes changes in activity between timesteps,

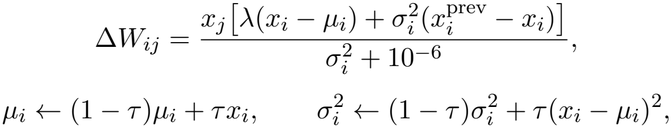

where the variance update uses the previous *µ*_*i*_ and 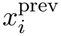 is the activity at the previous timestep. The Hebbian and predictive terms are weighted equally (*λ* = 1).

#### Reward prediction error rule (RPE).^**1**,**13**^

Hebbian coactivity is gated by a global scalar error *δ* computed at the input layer,

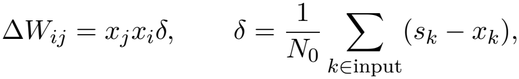

where *N*_0_ is the number of input neurons. It is the only rule with a non-local factor.

#### Common form

Writing each rule as Δ*W*_*ij*_ = *x*_*i*_*x*_*j*_*ε*_*ij*_ (Eq. 1; Table 1) gives

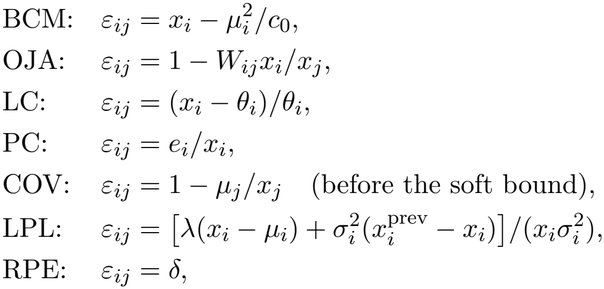

where the division is for exposition only and is never computed. OJA, PC and RPE have three hyperparameters (*α, β, η*); BCM, LC, COV and LPL have a fourth, *τ*. *c*_0_, *w*_max_, *ρ* and *λ* are fixed at the values above.

### 4.3 Environments

In every adaptive-behavior environment the environment’s only output is the scalar *s*_*t*_ ≥ 0 delivered to the input neuron. There is no separate reward channel: the same input drives activity, gates plasticity (through the presynaptic factor *x*_*j*_) and, for RPE, sets *δ*.

#### Deterministic bandit

One arm (a) always returns *s* = *r* (reward magnitude; *r* = 1 unless varied) and the other (b) returns 0. Performance is P(correct), the fraction of choices of the rewarded arm.

#### Two-armed bandits

In the unbaited bandit, the chosen arm *k* returns *s* ~ Bernoulli(*p*_*k*_), drawn afresh every trial, so always choosing the richer arm is optimal. In the baited bandit, a reward stays on an arm until that arm is chosen, and empty arms are re-baited each step with probability *p*_*a*_ or *p*_*b*_, so occasionally sampling the poorer arm is good and matching is near-optimal. ^17^

#### Inter-trial intervals (ITIs)

In some environments *k* null steps (*s* = 0, choices ignored) separate successive decisions (ITI = *k*). The network keeps updating during null steps, and all behavioral statistics use decision steps only.

#### Block schedules

Where contingencies change within a session (Fig. 5d–f, Fig. 3d and Fig. 4i– k), the network state (activities, weights, running statistics) carries across block boundaries, and there is no cue given to networks to mark a switch.

#### Matching pennies

The computer opponent predicts the agent’s next choice from its history. Using a buffer of the last 100 choices, it finds every earlier occurrence of the agent’s most recent three choices and tallies the choice that followed. If a two-sided *z*-test against 0.5 is significant (*p <* 0.05), it predicts the majority continuation; otherwise it guesses at random. The agent receives *s* = 1 when its choice differs from the prediction. In the non-competitive condition both arms pay with probability 0.5 regardless of history (unbaited). Sessions have seven 20,000-step phases: competitive, then three cycles of non-competitive and competitive (see Costly stochasticity, below). The network state is carried across phases and the opponent’s buffer is reset at each phase.

#### Feature selectivity

Following Law & Cooper, ^29^ the stimulus bank has 15 rectified, sharpened sinusoids of length 37, *f* (*n*) = max 0, sin(2*πn*/37) ^20^, circularly shifted to 15 evenly spaced phases. One pattern is drawn uniformly at random each step.

#### Slow features

Following Halvagal & Zenke, ^31^ the stimulus is two-dimensional, s_*t*_ = (*z*_*t*_, |*ξ*_*t*_|), with a binary slow channel *z*_*t*_ that switches with probability 0.05 per step and a fast channel *ξ*_*t*_ ~ N(0, *σ*^2^), *σ*^2^ ∈ {0.1, 10}.

### 4.4 Parameter sweeps and operating points

#### Grid

For every rule and both architectures we swept *α* and *β* over 15 log-spaced values in [10^−5^, 1], *η* over 6 log-spaced values in [10^−6^, 10^−1^] and, for BCM, LC, COV and LPL, *τ* over 4 log-spaced values in [10^−3^, 10^−1^] (1,350 or 5,400 cells per rule and architecture). Each cell was simulated with 20 independent networks (seeds), vectorized with JAX `vmap` and distributed over SLURM array jobs. Session lengths were 100,000 steps (deterministic bandit), 50,000 per reward condition (bandits), 20,000 per phase (matching pennies), 50,000 per noise level (slow features) and 10,000 (feature selectivity).

#### Task scores

All scores are averaged over seeds (Fig. S1–S3).

##### Reward seeking

P(correct) in the deterministic bandit over the second half of the session.

##### Probability matching

Seven conditions with (*p*_*a*_, *p*_*b*_) = (0.05*k*, 0.3 − 0.05*k*), *k* = 0, …, 6, each simulated separately. From the second half of each session we take the choice ratio *c*_*k*_ (fraction of choices of arm b) and compare it with the matching target *m*_*k*_ = *p*_*b*_/(*p*_*a*_ + *p*_*b*_), which runs linearly from 1 to 0. The signed matching index is

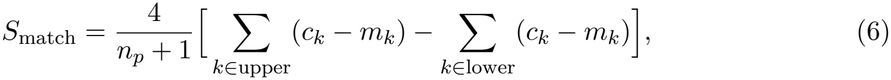

where “upper” is the three conditions with *m*_*k*_ *>* 0.5, “lower” the three with *m*_*k*_ *<* 0.5, and *n*_*p*_ = 7. *S*_match_ ≈ 0 is matching, *S*_match_ *>* 0 is more extreme than matching and *S*_match_ *<* 0 is flatter. The index was computed for both the baited and the unbaited bandit. We call a rule probability-matching in a task if its best cell has |*S*_match_| ≤ 0.35. The large network was swept only in the baited bandit, so for the unbaited bandit we simulated each rule at its best baited cell (20 seeds, same protocol); the classification was unchanged: PC (−0.25), RPE (−0.01), COV (0.00), LPL (−0.03) and OJA (−0.09) matched, and BCM (−0.60) and LC (−0.48) did not.

##### Costly stochasticity

3-bit block entropy 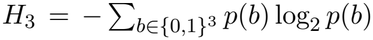 of the choice sequence over the second half of each matching-pennies phase. The sweep heatmaps (Fig. S2–S3) show the drop in median entropy from the competitive to the non-competitive phases over the settled phases (3–7).

##### Heavy-tailed exploration

Run lengths (consecutive identical choices) from the deterministicbandit sessions were pooled over seeds and fitted with a discrete power law (*l*_min_ = 1; see *Run-length distributions*). The score is −|*γ*− 2|, where *γ* is the fitted exponent; cells with fewer than 30 runs were skipped.

##### Feature selectivity

At the end of the session, the response *y*_*k*_ of the output neuron to each pattern *k* was computed from rest with three noise-free feedforward passes and rectification. The score is 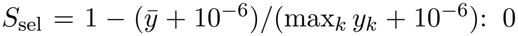 for uniform responses, approaching 1 when a single pattern dominates.

##### Slow-feature selectivity

*χ* = ⟨*y*⟩ _*z*=1_ ⟨*y*⟩ _*z*=0_ /(max_*t*_ *y*_*t*_ min_*t*_ *y*_*t*_ + 10^−6^) over the last 1,000 steps. The score is the minimum over the two noise levels.

For the behavior grid (Fig. S1) each rule is shown at the cell that maximizes each score over the whole grid; for probability matching and heavy tails this is the cell with the index closest to 0 or the exponent closest to 2. The heatmaps (Fig. S2–S3) show the best value over *η* and *τ* at each (*α, β*).

#### Reward-seeking operating point

Unless stated otherwise, each rule and architecture is simulated at its best deterministic-bandit cell: the cell that maximizes the median over steps 80,000–100,000 of the across-seed mean P(correct), with *η* ≥ 10^−5^. Averaging across seeds and excluding the lowest noise level avoids selecting cells in which the network stays frozen near its initialization. Figure-specific departures from this operating point, and the reasons for them, are given below and in Table S2. For BCM and LC in the large network, the unrestricted best cell had *α* = 1 and *β* = 10^−5^, where networks chose the rewarded arm equally well with plasticity switched off; for these two we excluded *α* = 1 and *β* = 10^−5^ (Section S1).

### 4.5 Signal seeking (Fig. 1)

Fig. 1b,c shows a single network with *α* = *β* = 0.01 and *η* = 0.005, driven with *s* = 1 for 500 steps and then *s* = 0 for 500 steps, with activities initialized to *x*_in_ = 0 and *x*_out_ = 1. For Fig. 1e–g, each rule and architecture was re-simulated at its operating point for 100,000 steps with 50 seeds. Learning curves are per-seed means in consecutive 100-step bins, summarized as the median and interquartile range across seeds. Fig. 1g reports the operating-point score itself: the median over steps 80,000–100,000 of P(correct) averaged across the 50 seeds, with 95% bootstrap confidence intervals over seeds (1,000 resamples).

### 4.6 Reward magnitude (Fig. 2, Fig. S4)

#### Rate and asymptote sensitivity

For each rule, at its large-network operating point, we simulated the deterministic bandit at reward magnitudes *r* ∈{0.2, 0.4, 0.6, 0.8, 1.0} (50 seeds, 100,000 steps). Per-seed choices were binned in 100-step windows and the median across seeds was taken at each bin. The asymptote *A*(*r*) is the mean of this curve over the final 10% of bins. Time to criterion *T* (*r*) is the first bin at which the curve reaches 0.9 *A*(*r*). The criterion is relative to each level’s own asymptote, so a larger reward is not credited with faster learning merely because its curve plateaus higher, as it would be under a fixed threshold, and no level is censored. Rate is 1/*T* (*r*). Rate sensitivity and asymptote sensitivity are the least-squares slopes of rate and of *A*(*r*) against *r* (at least three usable levels required). Rate sensitivity is displayed on a symmetric-log axis. Animal-like behavior, in which larger rewards speed learning without raising the plateau, ^32^ corresponds to positive rate sensitivity with near-zero asymptote sensitivity (Section S2).

#### Sample efficiency (Fig. 2d)

PC (large network) was simulated at ten reward magnitudes *r* ∈ {0.1, …, 1.0} for 20,000 steps, in 20 independent batches of 20 seeds. For each batch, time to criterion was the first 100-step bin at which the across-seed median of the binned curve reached P(correct) = 0.9 (20,000 if never reached). Points are batches; bars are mean ± s.e.m. across batches. The learning curves (right) use 300 seeds at *r* ∈ {0.2, 0.5, 1.0}.

#### Robustness across parameters (Fig. S4)

The same measurement was repeated at every signal-seeking cell of each rule and architecture (12,880 cells; 20 seeds; activities capped at 3); cell selection and simulation budget are in Section S2.

#### Mouse data (Fig. 2b)

The left panel is our reanalysis of the hidden-target navigation data of Gong et al. ^32^ (figshare 10.25378/janelia.32145868; 5, 10, 20, 50 and 100 µL groups, *n* = 14, 5, 5, 5, 14 mice). For each mouse, the sequence of target latencies on successful trajectories, concatenated across training, was smoothed with a causal 50-trajectory moving mean. With the initial latency *B*_0_ taken as the median of the first 15% of the smoothed curve and the trained latency *A* as the median of the last 10%, reinforcements to criterion is the first trajectory at which the smoothed latency falls below *A* + 0.1(*B*_0_ − *A*), that is, 90% of the way to trained performance. The authors’ exact criterion is not in their public code, so this is a reconstruction; it reproduces the published rank order across reward sizes (group means 586, 178, 89, 70 and 52 reinforcements). The right panel reproduces the joystick learning curves of Gong et al. (their Fig. 4B), digitized from the published figure because those data are not public.

### 4.7 Probability matching (Fig. 3, Fig. S5)

#### Animal data

We reanalyzed mouse choices from the dynamic-foraging task of Bari et al. ^51^ (baited; blocks with 4:1 or 8:1 reward-probability ratios; 585 sessions, 204,916 trials; one animal with incomplete data excluded, and the final block of each session dropped because its probabilities were not recorded) and from the two-armed bandit of Beron et al. ^35^ (unbaited; high/low probabilities of 0.9/0.1, 0.8/0.2 or 0.7/0.3; 525 sessions, 378,351 trials; blocks delimited by the task’s trial counter). The reward ratio of a block is *p*_*A*_/(*p*_*A*_ + *p*_*B*_) and the choice ratio is the fraction of choices of side A. Choice ratios were computed per block after dropping the first 10 trials, from blocks longer than 50 (Bari) or 20 (Beron) trials, and are shown as mean ± s.d. across blocks. The s.d. of the block choice ratio was computed from all trials of blocks longer than 50 trials.

#### Model fit

A small PC network was run in one continuous unbaited-bandit session of 60 blocks with the Beron conditions, with probabilities scaled by 0.3 so that the total reward rate per trial was 0.3 (for example, 0.27/0.03 for the 90:10 condition). Block lengths were geometric with mean 5,000 trials (minimum 10), and conditions were visited in balanced random order. Twenty seeds shared the same block schedule, with each seed treated as a session. Parameters were chosen to minimize the mean squared deviation of each block’s choice ratio from the Beron choice-ratio curve, interpolated at the block’s reward ratio, computed from blocks longer than 2,600 trials after discarding the first 2,500 trials. Targets were Beron per-condition means over blocks longer than 50 trials. One-dimensional scans of each parameter, followed by hand adjustment of *α* and *β* together around the scan optima, gave *α* = 0.023, *β* = 0.15, *η* = 0.005, ITI = 0 and reward scale 1.0 (Fig. S5). Model choice ratios use the same block filter and skip. The model’s choice-ratio s.d. uses all trials of blocks longer than 2,600 trials.

### 4.8 Costly stochasticity (Fig. 3h–k)

#### Animal data

Mouse choices are from Aloor et al. ^36^ (G-Node GIN release). We used the seven mice that alternated matching-pennies sessions with sessions against a random opponent (coin flip, “Algorithm 0”): four behavioural mice and three widefield-imaging mice, each with 5–6 coin-flip sessions. For each mouse we used its coin-flip sessions and the matching-pennies sessions interleaved with and immediately bracketing them, dropped missed trials and sessions with fewer than 100 trials, and averaged the 3-bit entropy over sessions of each type.

#### Protocol and parameter choice

Each rule was swept in the small network over the grid above with seven 20,000-step phases (competitive, then three cycles of non-competitive and competitive; 20 seeds). Phases 3–7 were treated as settled. Cells were kept if behavior had settled, meaning that repeated phases of the same type differed by less than 0.1 bits in median entropy, and if choices were unbiased against the computer (median side bias | *P* (choice A) − 0.5| *<* 0.05). Among these, we took the cell with the largest drop in median entropy from the competitive to the non-competitive phases: BCM (*η* = 10^−4^, *τ* = 10^−3^, *α* = 0.037, *β* = 1), OJA (10^−2^, *α* = *β* = 0.0014), LC (10^−6^, *τ* = 0.022, *α* = 0.44, *β* = 0.085), PC (10^−3^, *α* = 0.016, *β* = 0.44), COV (10^−6^, *τ* = 10^−3^, *α* = 0.0032, *β* = 0.037), LPL (0.1, *τ* = 0.1, *α* = *β* = 1) and RPE (10^−6^, *α* = 0.085, *β* = 1), with *η* given first. Each chosen cell was then re-run with 50 new seeds. In the sweep each phase restarted the noise sequence from the same seed; in the re-runs the random state is carried across phases, so every phase receives new noise and opponent draws. The two give the same settled values.

#### Measures

Choice entropy is the 3-bit block entropy (Task scores, above), computed for each seed over the second half of each settled phase and averaged across phases of the same type (Fig. 3j). For the traces in Fig. 3i, entropy was computed in a trailing window of 30 trials for mice and 300 trials for the model, which is shown over 10 times as many trials. The model example seed is the one closest to the median entropy in both conditions. Its competitive segment is the last 3,000 trials of the final phase. Its non-competitive segment is, among its 3,000-trial segments in the second halves of the non-competitive phases whose preferred side changed at least once (in 300-trial blocks; 57% of all seeds’ segments do), the one whose mean windowed entropy is closest to the median over all seeds. Side bias is |*P* (choice A) −0.5|, and P(switch) is the fraction of trials on which the choice differs from the previous one.

### 4.9 Explore–exploit states (Fig. 4a–c, Fig. S6)

We fitted two-state hidden Markov models (HMMs) with Bernoulli emissions to the rewarded/ unrewarded choice sequences of the small-network deterministic-bandit sweep (last 50,000 steps of each session; 20 seeds per cell, sharing parameters). For each rule we took the three *η* values with the most cells above 0.7 reward (for rules with *τ*, at the *τ* with the most such cells) and fitted every cell above 0.7 at those *η* values (892 cells; BCM and OJA had none). Fits used scaled Baum–Welch EM run to a relative log-likelihood change below 10^−6^, with state 1 (“exploit”) defined as the state with the higher emission probability. States were decoded with the Viterbi algorithm. A cell was called discretely bistable if 2Δ log *L >* 200 relative to a one-state model, *p*_1_ − *p*_0_ *>* 0.25, the median number of state switches per seed was at least 3, and the mean dwell time in each state was at least 1.5 trials. For each rule the representative cell (Fig. 4b,c) is the bistable cell with the largest minimum switch count across seeds. Recurrence was confirmed by re-simulating these cells for 5 × 10^6^ steps with 10 new seeds (Fig. S6b).

### 4.10 Run-length distributions (Fig. 4d–h)

A small PC network (*α* = 0.0373, *β* = 0.0072, *η* = 0.01, the forgetting-task parameters; weights initialized with *κ* = 0.25; activities capped at 3) was run on a baited bandit at ITI = 0 or 2, with reward probabilities (1, 0), (0.45, 0.15) or (0.05, 0.05) (24 seeds, 4,000 burn-in trials, then 120,000 trials). Run lengths (consecutive identical choices on decision steps) were pooled across seeds that chose the better arm on more than half of trials (all seeds for symmetric arms). Distributions were fitted following Clauset, Shalizi & Newman ^58^ with the `powerlaw` package (Alstott et al. 2014^102^): a discrete power law with *l*_min_ = 1, fitted by maximum likelihood, was compared with an exponential by the normalized log-likelihood ratio (Vuong test). A distribution was called a power law or exponential if *p <* 0.05, and not significant otherwise. The example runs in Fig. 4e–h use the same network with *η* = 0.002 (reward probabilities 0.45/0.15; ITI = 0, seed 6; ITI = 2, seed 8; 360,000 trials). The lower noise makes the attractor geometry visible without changing the power-law or exponential verdicts. The first-passage account of heavy tails in the main text is only an approximation, because network activity also decays toward zero; this may explain why some fitted exponents in Fig. S2–S3 are more negative than expected for a pure random walk, *P* (*l*) ∝ *l*^−3/2^.

### 4.11 Interference-dependent forgetting (Fig. 5a,b, Fig. S7)

Small networks (250 seeds) were trained for 100,000 steps on the deterministic bandit and then, from the same post-training state, run for 100,000 steps under each of three conditions: no input (both arms return 0), constant input (both arms return 1) and reversal (the previously unrewarded arm returns 1). The stored memory is *W*_diff_ = *w*_*a*_ − *w*_*b*_ (with *w*_*k*_ = *W*_*k*,in_), where a is the originally rewarded arm; curves show mean ± s.d. across seeds. Several rules (PC, COV, LPL, RPE) reach the maximal deterministic-bandit score over a plateau of tied cells, and the lowest-noise cell on such a plateau barely moves under any condition. For these rules we searched the plateau for the cell with the strongest contrast between no-input and constant-input retention (Section S1). Statistics are described under *Statistics*.

### 4.12 Memory retention across alternating contexts (Fig. 5d–f)

Networks with two input and two output neurons were run through 10 blocks of 50,000 steps (100 seeds). The rewarded output alternated between environments A and B (output a rewarded in A blocks, output b in B blocks), and reward was delivered to input neuron 1 during A blocks and to input neuron 2 during B blocks, so each context has its own synapses. The input when rewarded was *s* = 1 on the active context’s input neuron. Each rule used its forgetting-task parameters (Table S2). The preference of each channel is the difference of its two outgoing weights, signed so that positive means correct: 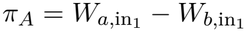 and 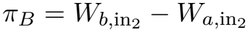. To compare rules whose weights live on very different scales, preferences are divided by each seed’s trained scale: the mean absolute preference of the active channel at the end of blocks 2–10. Seeds whose trained scale was below 0.005 (no learning) or above 1 (divergence) were excluded, and the per-rule *n* is given in the legend. Fig. 5f shows the current and inactive channels’ normalized preferences at the end of block 10 (box plots: median, interquartile range and 5th– 95th percentiles across seeds). Fig. 5e shows the raw preferences over the first five blocks as the median and 35th–65th percentile band across seeds. The summary of held memory versus accuracy (Fig. S8) is defined in Section S3.

### 4.13 Attractor analyses

All attractor analyses use PC in the small network, in the reduced coordinates *W*_diff_ = *w*_*a*_− *w*_*b*_ and *X*_diff_ = *x*_*a*_ − *x*_*b*_ together with *x*_in_ (Li & Erlich ^48^). Subtracting the activity and PC weight updates (Eq. 3 and Eq. 5) of the two output neurons gives a closed system in these coordinates. Its fixed points form the line *X*_diff_ = *s W*_diff_, which is marginally stable along *W*_diff_.

#### Phase portraits (Fig. 1j)

Deterministic flows of the reduced system were integrated for *s* = 1 and *s* = 0 (*α* = *β* = 0.01). Stochastic trajectories of the full model on the deterministic bandit (*α* = 0.02, *β* = 0.03, *η* = 0.008, activities capped at 3; 2,000-step burn-in, 40,000 steps shown; three seeds) show both branches.

#### Reward magnitude (Fig. 2e–i)

PC was run at *α* = 0.0848, *β* = 0.0164 and *η* = 0.003, with activities not capped. (*α, β*) is one of the top-scoring small-network deterministic-bandit cells at *η* = 10^−2^, and *η* was lowered so that activity noise is small compared with the output difference a learned weight produces at the smallest reward (Section S1). Learning curves (Fig. 2f) are across-seed means (10 batches of 80 seeds, 300,000 steps) at *r* ∈{0.2, 0.3, 0.5, 0.7, 1.0}. Separate runs of 200 seeds (300,000 steps each) were used for the attractor analyses. Attractor slopes (Fig. 2g) are through-origin fits of *X*_diff_ on *W*_diff_ in the central *x*_in_ band over the last 50,000 steps. Drift *v*(*W*_diff_) and diffusion *D*(*W*_diff_) (Fig. 2h) were estimated from binned increments at a lag of 400 steps from 120 of these seeds. Learning curves were then predicted with no fitted parameters for the 80 held-out seeds by integrating the Fokker–Planck equation 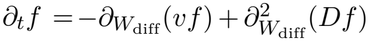, where *f* (*W*_diff_, *t*) is the density of *W*_diff_ across networks, forward from their initial *W*_diff_ distribution, using an upwind finite-volume scheme (Fig. 2i). Full details and validation are in Li & Erlich. ^48^

#### Forgetting (Fig. 5c)

Networks with the PC forgetting-task parameters (120 seeds) were trained for 100,000 steps and then run for 20,000 steps under each retention condition, or with continued training as a reference. Traces show *W*_diff_ for 50 of the 120 seeds over the last 10,000 training steps and the 20,000 retention steps. Clouds show the last 1,200 steps; one example trajectory is overlaid.

#### Attractor length (Fig. 3e–g)

PC (*α* = 0.01, *η* = 0.005, activities capped at 3) was run on a baited bandit with total reward rate 0.45, (*p*_*a*_, *p*_*b*_) = 0.45 (*q*, 1 − *q*), for nine reward ratios *q* ∈ [0.1, 0.9] (40 seeds; 40,000-step burn-in; 40,000 steps sampled). The weight rate *β* sets attractor length (short attractor, *β* = 0.02; long attractor, *β* = 0.15; sweep *β* ∈ {0.02, 0.03, 0.04, 0.05, 0.1, 0.15}). Within-block choice s.d. is the s.d. of the choice ratio across 20 equal windows of the sampled period, averaged over seeds. Attractor length *L*/*W* was measured at *q* = 0.5: the cloud in the (*X*_diff_, *W*_diff_) plane was split into its two branches by the sign of *W*_diff_, and each branch was rotated onto its principal axes, with *L* and *W* the 1st–99th-percentile extents along the first and second axes, averaged over branches weighted by occupancy.

#### Costly stochasticity (Fig. 3k)

PC at its matching-pennies cell (above; 50 seeds). Clouds show *W*_diff_ against *X*_diff_ over the last 5,000 steps of the final non-competitive phase and the final competitive phase.

#### Changing environments (Fig. 4i–k)

One small PC network per seed (*α* = 0.1, *β* = 0.01, *η* = 0.005, *κ* = 0.25, activities capped at 3; 48 seeds) was run through five 20,000-step blocks with the state carried across: deterministic (1, 0), deterministic (0, 0.4), unbaited bandit (0.15, 0.45) with ITI = 1, constant input with no feedback (1, 1), and deterministic (0, 0.4) again (values for arms a, b). Choice traces are 400-step moving averages (median and IQR across seeds). Run lengths pooled across seeds within each block were fitted as above. Clouds use the second half of each block from six seeds.

### 4.14 Statistics

The unit of analysis is the seed (an independently initialized network); timesteps within a seed are never treated as independent. For forgetting (Fig. S7, Table S1), each seed’s retention is the mean *W*_diff_ over the last 20% of the retention phase, also reported as a fraction of that seed’s trained *W*_diff_ (mean over the last 20% of training). No-input and constant-input retention (paired, same trained state) were compared with one-sided Wilcoxon signed-rank tests, and reversal retention was tested against zero with one-sided one-sample Wilcoxon tests. Paired and one-sample *t*-tests were run as robustness checks, and effect sizes are matched-pairs rank-biserial correlations. *p*-values were Benjamini–Hochberg corrected across the 14 tests (7 × rules 2 questions). For memory retention (Fig. 5f), a rule retained the inactive context if at least 95% of its seeds had an inactive-context preference above zero. This was tested with exact one-sided binomial tests (*H*_0_: fraction ≤ 0.95), Holm–Bonferroni corrected across the rules with at least 50 usable seeds (PC, COV, LPL and RPE). For matching pennies (Fig. 3j), settled-phase entropies against the computer and in the non-competitive phases were compared with one-sided paired Wilcoxon tests (50 seeds per rule), Holm corrected across rules (competitive higher in 50 of 50 seeds for six rules and 49 of 50 for OJA). For mice, entropy against the computer and in coin-flip sessions was compared with a one-sided paired Wilcoxon test across mice (*n* = 7). Power-law comparisons used the Vuong likelihood-ratio test described above. All tests are two-sided unless stated.

### 4.15 Software, data and code availability

Simulations were written in Python with JAX^103^ (JIT-compiled and vectorized) and run on the Sainsbury Wellcome Centre HPC cluster via SLURM. Analyses used NumPy,^104^ SciPy,^105^ pandas ^106^ and powerlaw,^102^ and figures were made with Matplotlib ^107^ and Adobe Illustrator. Code and the notebooks that generate each figure are available at https://github.com/ccli3896/hebbian-flexible-attractors. The code is currently released with all rights reserved, and we intend to release it under an open-source license on publication. Simulation results (parameter sweeps) are available on Zenodo (doi:10.5281/zenodo.24342269). The mouse data are from Aloor et al. ^36^ (matching pennies; G-Node GIN, https://gin.g-node.org/DELab/matching-pennies-widefield-2026), Bari et al. ^51^ (dynamic foraging; obtained from the authors), Beron et al. ^35^ (two-armed bandit; publicly available on Harvard Dataverse, doi:10.7910/DVN/7E0NM5) and Gong et al. ^32^ (hidden-target navigation; figshare, doi:10.25378/janelia.32145868).

## Acknowledgements

We thank the Sainsbury Wellcome Centre Scientific Computing team for access to the HPC cluster.

## Author contributions

Contributions are given using the CRediT taxonomy. C.L.: Conceptualization, Methodology, Software, Formal analysis, Investigation, Visualization, Writing – original draft, Writing – review & editing. J.B.: Formal analysis and Investigation (theory and simulations for an earlier version of this work), Writing – review & editing. H.Y.: Formal analysis (theory of explore/exploit run-length distributions), Visualization, Writing – review & editing. B.D.P.: Formal analysis (theory of explore/exploit run-length distributions), Writing – review & editing. M.K.: Investigation (preliminary work on matching pennies). J.A.: Resources (matching-pennies mouse data). C.A.D.: Resources (matching-pennies mouse data), Funding aquisition. J.C.: Funding acquisition, contributed to discussions. J.C.E.: Conceptualization, Supervision, Funding acquisition, Writing – review & editing.

## Use of AI

An AI assistant (Claude, Anthropic) was used to help write code for the parameter sweeps and to proofread and organize the text. The manuscript was written by the authors, who designed and checked all analyses and take full responsibility for the content.

