## Supplementary Figures and Methods for "Adaptive behaviors emerge from Hebbian-like plasticity through flexible attractors"

### Supplementary Information for "Adaptive behaviors emerge from Hebbian-like plasticity through flexible attractors"

This file contains Supplementary Figures S1–S8, Supplementary Methods S1–S3 and Supplementary Tables S1–S2.

#### Contents

**Supplementary Figures** 2

**Supplementary Methods** 10

**S1 Parameter selection** 10

**S2 Reward-magnitude sweep** 10

**S3 Memory-retention summary** 11

**Supplementary Tables** 12

#### Supplementary Figures

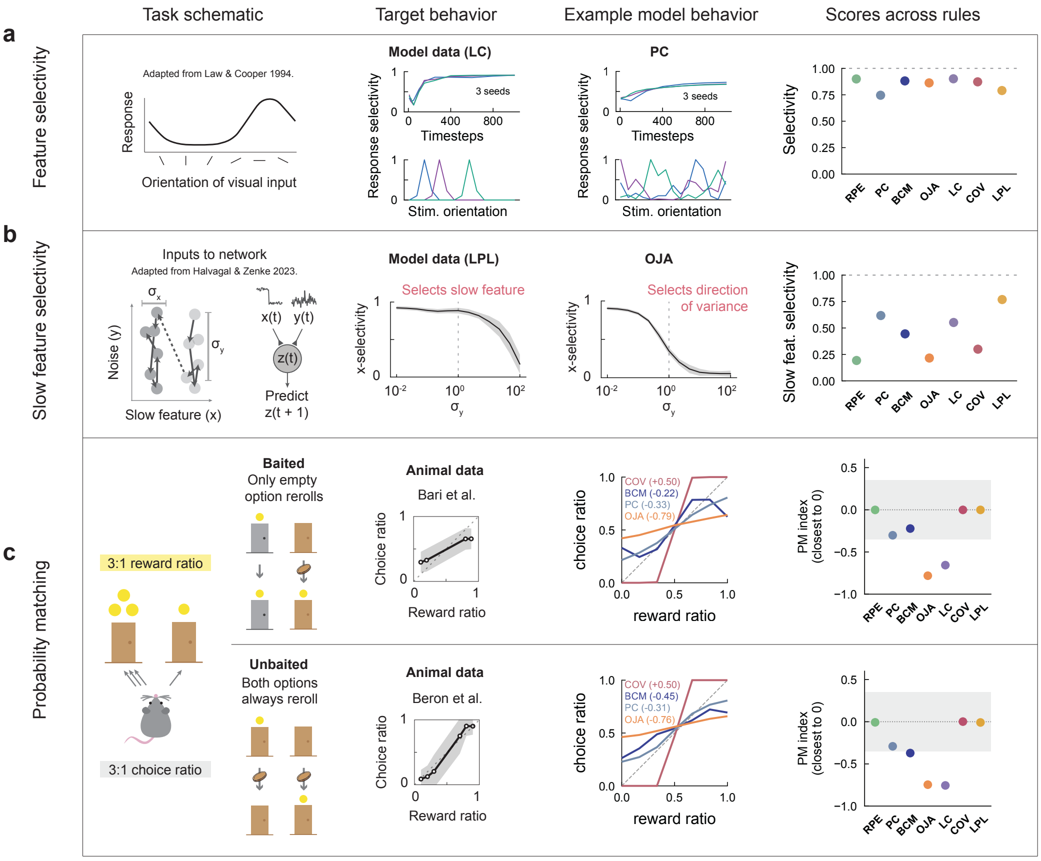


**Supplementary Figure S1: Behavior grid: representation learning and probability matching across rules.** Tasks whose comparison across all seven rules is not shown in the main figures (reward seeking across rules is in Fig. 1g, costly stochasticity in Fig. 3h–k). **a**, Feature selectivity. Schematic: orientation-tuned response profile (adapted from Law & Cooper[1]). Target behavior: model response selectivity over training and across stimulus orientation, LC rule, 3 seeds. Example model behavior: same for PC. Scores: selectivity (0–1) across all seven rules.
**b**, Slow-feature selectivity. Schematic: adapted from Halvagal & Zenke.[2] The network receives a slow, informative feature $x(t)$ mixed with fast noise $y(t)$ and predicts $z(t+1)$; $x$-selectivity is scored across noise level $\sigma_y$. Target behavior: LPL, which selects the slow feature ($x$-selectivity $\to 1$ as $\sigma_y$ grows). Example model behavior: OJA, which instead selects the direction of highest variance regardless of timescale ($x$-selectivity $\to 0$ as $\sigma_y$ grows). Scores: slow-feature selectivity (0–1) across all seven rules. Only a subset of rules solve for slowness rather than raw variance.
**c**, Probability matching. Schematic: two-armed bandit at a 3:1 reward ratio, baited (unclaimed reward persists; only the empty arm rerolls) vs. unbaited (both arms reroll every trial). Target behavior: animal data, baited (Bari et al.[3]) and unbaited (Beron et al.[4]), choice ratio vs. reward ratio. Example model behavior: same axes for four rules (COV, BCM, PC, OJA), each at the cell with its highest combined baited and unbaited matching index, annotated with that index ($S_{\mathrm{match}}$; positive, steeper than matching, toward always choosing the richer arm; negative, flatter, toward indifference). Scores: for each rule, the matching index closest to 0 (0, perfect matching; shaded band, $|S_{\mathrm{match}}| \leq 0.35$, the criterion in Methods), shown separately for baited and unbaited.

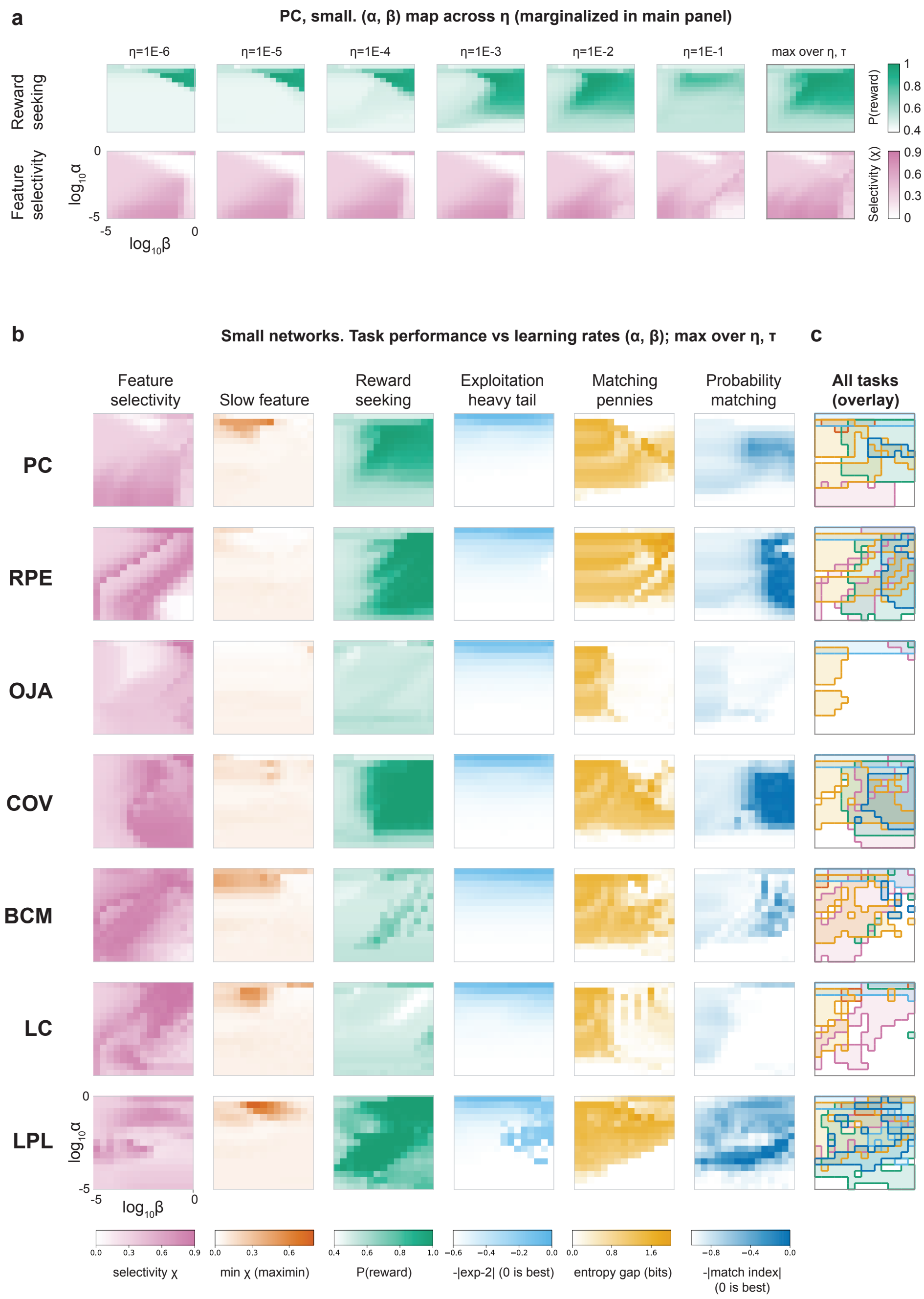


**Supplementary Figure S2: Parameter sweeps, small network.** **a**, Marginalization example (PC). Illustrates what the $\alpha \times \beta$ maps marginalize over: for the PC rule, reward seeking (top row, green, P(reward)) and feature selectivity (bottom row, magenta, selectivity $\chi$) are shown as $\alpha \times \beta$ heatmaps at each of the six swept $\eta$ values ($\tau$ collapsed within each panel by taking the maximum, for applicable rules), plus a rightmost "max over $\eta$, $\tau$" envelope, which is the single per-$(\alpha, \beta)$ best achievable value used throughout panel **b**.

**b**, Performance grid, small networks (no hidden layer). Rows are the seven plasticity rules; the first six columns are task-specific $\alpha \times \beta$ heatmaps (each cell $=$ max over $\eta$, $\tau$, i.e. the best achievable performance at that learning-rate pair, for visualization purposes); the seventh column overlays all six tasks' "good-performance" regions for that rule. Each task uses a single identity hue (white, worst; saturated, best) shared between its heatmap and its overlay colour: feature selectivity (magenta, selectivity $\chi$, 0–0.9), slow feature (orange, maximin $\chi$ over two slow-probability conditions, 0–0.8), reward seeking (green, P(reward), 0.4–1.0), heavy tail (blue, $-|$Lévy exponent $- 2|$, 0 best), matching pennies (amber, entropy gap $=$ median entropy vs. computer opponent minus vs. random reward, settled phases, 0–2 bits) and probability matching (dark blue, $-|$matching index$|$, 0 best).

**c**, Per task, every $(\alpha, \beta)$ cell whose performance clears an absolute threshold (translucent fill and outline in that task's colour). A rule that cannot solve a task shows nothing in that colour anywhere in its overlay, so the overlay column reveals both which tasks each rule can solve and whether their good regions coincide in learning-rate space. Axes: $\log_{10}\beta$ ($x$), $\log_{10}\alpha$ ($y$), both spanning $-5$ to 0.

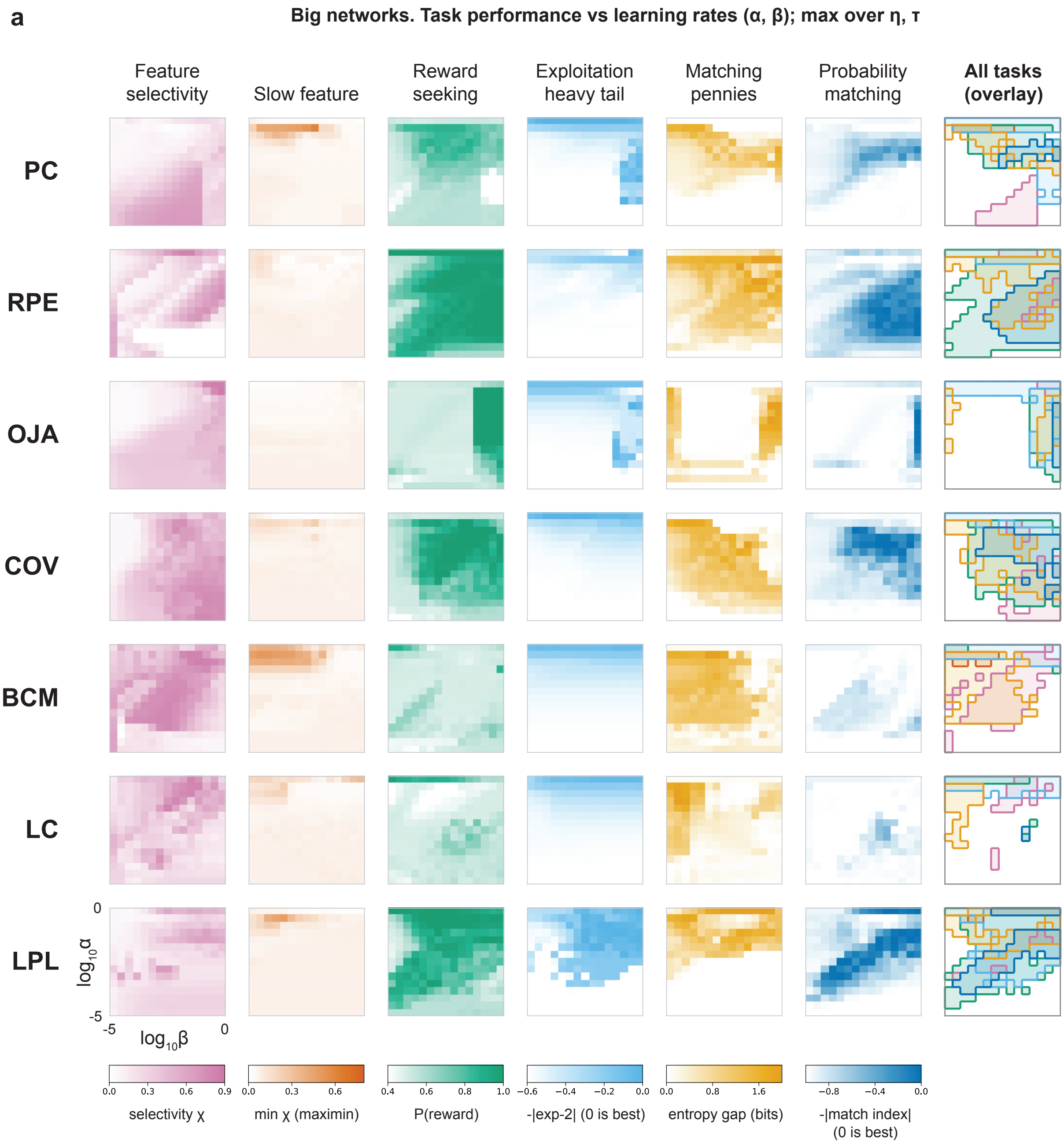


**Supplementary Figure S3: Parameter sweeps, large network.** Same layout, tasks, colour scale and overlay convention as Fig. S2b, computed instead for the large architecture (one hidden layer of 50 units). Several rules' good-performance regions shift at this larger width: for example, OJA's matching-pennies region splits into two disjoint high-entropy-gap bands rather than one contiguous region, and LPL's probability-matching region becomes a much larger, well-defined diagonal band. The overlay column again shows task-overlap patterns across rules, with differences from the small network most visible for RPE, LC and LPL.

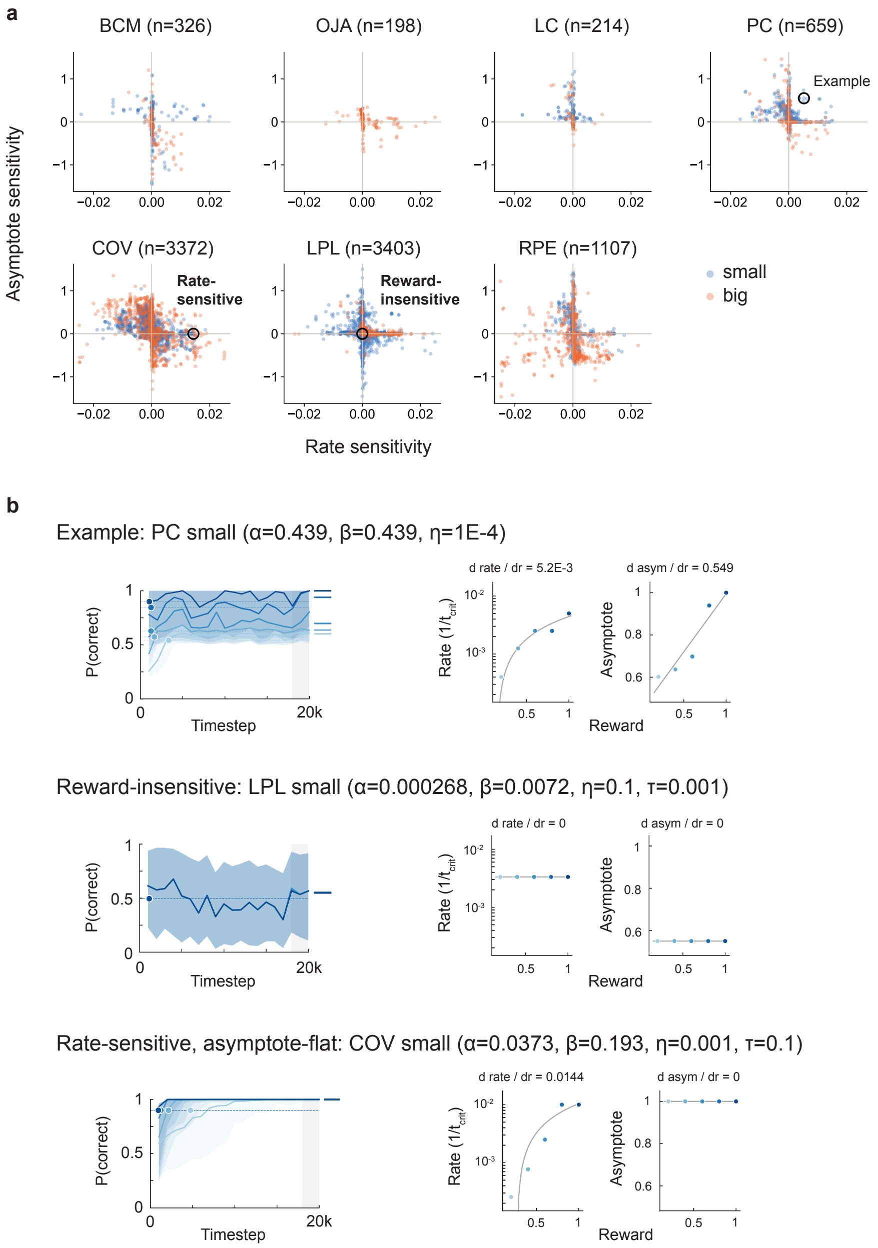


**Supplementary Figure S4: Reward magnitude: rate and asymptote sensitivity across parameters.** For every rule (BCM, OJA, LC, PC, COV, LPL, RPE) $\times$ architecture (small, large), hyperparameters were swept and each signal-seeking combination ("cell") was evaluated across five reward levels ($\eta$, $10^{-6}$–$10^{-1}$; $\alpha$, $\beta$, $10^{-5}$–$10^{0}$; $\tau$, $10^{-3}$–$10^{-1}$). Rate and asymptote sensitivities are the slopes $d(\mathrm{rate})/dR$ and $d(\mathrm{asymptote})/dR$ defined in Methods ($n = 9{,}279$ cells). In this sweep, activities were capped at 3 after each noise step and each cell was run with 20 seeds. Main-figure simulations (Fig. 2c) were re-run without the cap.

**a**, Population scatter. All rules span a range of asymptote and rate sensitivities. Each panel is one rule; points are individual hyperparameter cells at $(d(\mathrm{rate})/dR,\ d(\mathrm{asymptote})/dR)$, coloured by architecture (blue, small; orange, large). Axes are shared and centred at zero. The three example cells in **b** are ringed and labelled.

**b**, Individual examples. Three cells labelled in **a**: "Example" (strong positive sensitivity on both axes; PC small, $\alpha = 0.439$, $\beta = 0.439$, $\eta = 10^{-4}$), "Reward-insensitive" (both slopes $\approx 0$, LPL small, $\alpha = 0.000268$, $\beta = 0.0072$, $\eta = 0.1$, $\tau = 0.001$) and "Rate-sensitive, asymptote-flat" (COV small, $\alpha = 0.0373$, $\beta = 0.193$, $\eta = 0.001$, $\tau = 0.1$). Each is shown as three panels: learning curves per reward level (median $\pm$ IQR, criterion crossing marked, asymptote window ticked); rate vs. reward with linear fit (log-scaled); and asymptote vs. reward with linear fit.

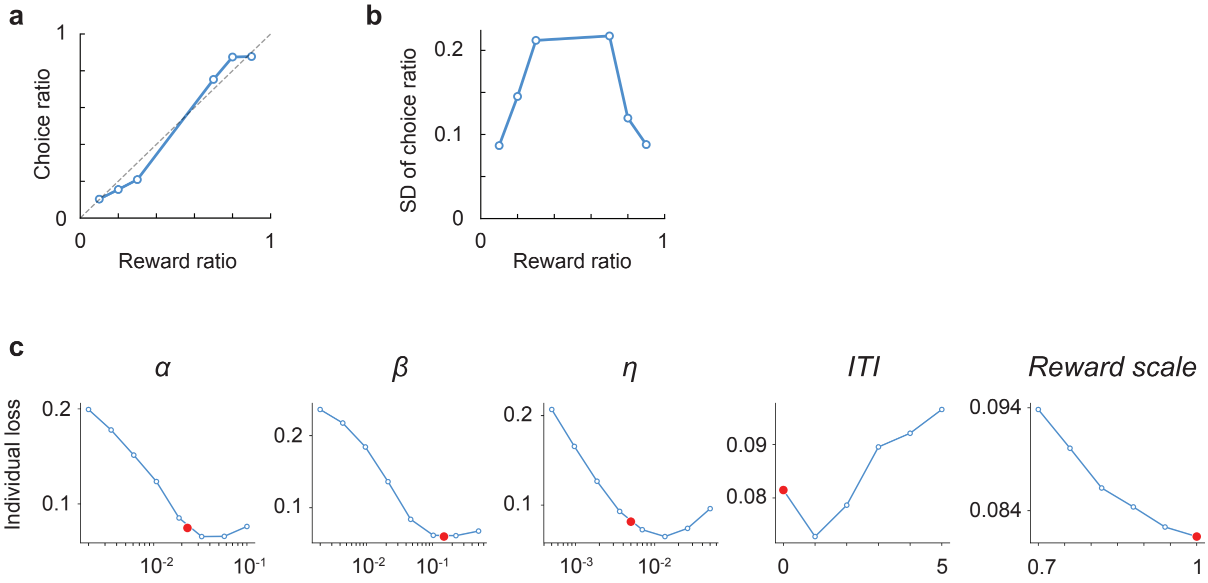


**Supplementary Figure S5: Probability matching: model fit to mouse data.** **a**, Fitting target: mean choice ratio vs. reward ratio for the Beron mice, from blocks longer than 50 trials using all trials (Fig. 3c uses blocks longer than 20 trials and drops the first 10).
**b**, Standard deviation of the block choice ratio vs. reward ratio (as in Fig. 3c, bottom right); shown for comparison and not used in the fit.
**c**, Sensitivity of the fit to each of the model's five parameters (activity rate $\alpha$, weight-update rate $\beta$, noise scale $\eta$, inter-trial-interval length ITI, reward scale), varied one at a time with the others held at $\alpha = 0.02$, $\beta = 0.05$, $\eta = 0.005$, ITI $= 0$ and reward scale 1.0. Red dots, the values used in Fig. 3d ($\alpha = 0.023$, $\beta = 0.15$, $\eta = 0.005$, ITI $= 0$, reward scale 1.0), each scored with the other parameters at the held values. $\alpha$ and $\beta$ were then adjusted together by hand around the slice optima, which also brought the model's s.d. of choice ratio closer to the animals'. $\eta$ and ITI are left at default values. Note that a range of parameter settings produce individual-block matching (Fig. S2–S3).

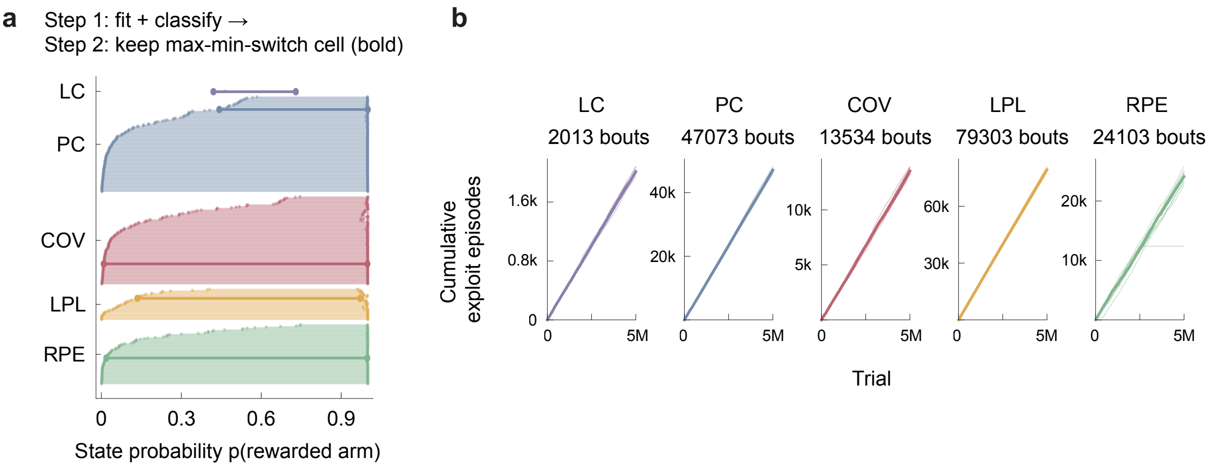


**Supplementary Figure S6: Explore–exploit states: HMM fits.** **a**, Two-state HMM fit to the rewarded/unrewarded choice sequence per swept cell. Cells were classified as discretely bistable using the criteria in Methods. The plot shows the fitted $(p_0, p_1)$ per discretely bistable cell, grouped by rule and sorted by $p_0$. Bistable/tested per rule: PC 166/184, COV 153/225, RPE 104/255, LPL 54/223, LC 2/5 (479/892 in total). The max–min-switch cell per rule (bold) is used in Fig. 4b,c and in **b**.
**b**, The same cells refit at $T = 5{,}000{,}000$ (10 new seeds, 100× longer). Cumulative exploit-episode count per seed; median in bold. All five climb linearly (median totals: LC 2,013; PC 47,073; COV 13,534; LPL 79,303; RPE 24,103). Refit $(p_0, p_1)$ (LC 0.422/0.730; PC 0.437/1.000; COV 0.009/0.999; LPL 0.201/0.972; RPE 0.020/0.998) closely matches **a** for 4 of 5 rules; LPL's $p_0$ shifts from 0.136 to 0.201. (In this rerun, LC and LPL used $\tau = 0.0251$ rather than their swept value, $\tau = 0.0215$.)

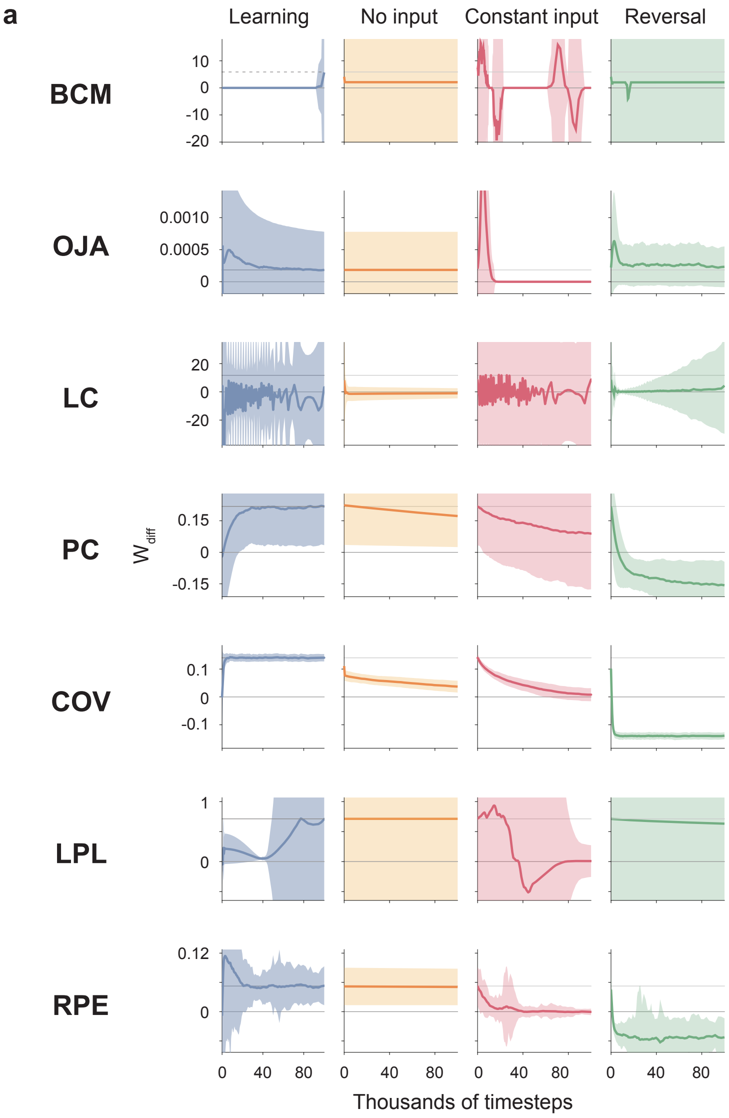


**Supplementary Figure S7: Interference-driven forgetting, all rules.** All learning rules on the tasks from Fig. 5a,b that test interference-driven forgetting. Rows are rules. Columns show learning (100k timesteps of training on the deterministic bandit), followed by a 100k-timestep retention phase under no input, constant input, or reversal; all three retention phases start from the same trained state. Memory is the weight difference $W_{\mathrm{diff}}$ between the originally rewarded and unrewarded arms. Lines are lightly smoothed means and shading is $\pm$ s.d. across 250 seeds; grey horizontal lines mark 0 and the trained level. Because weight magnitudes differ by orders of magnitude between rules, $y$-axes are scaled per rule to the range of the mean lines, and the s.d. bands of the most variable rules (BCM, LC and LPL) extend beyond the axis limits. See Table S1 for statistics and comparisons. Networks used the best-performing hyperparameters on the deterministic task for each rule (Fig. 1g) and were then filtered for performance on the forgetting task (Methods; Section S1; parameters in Table S2).

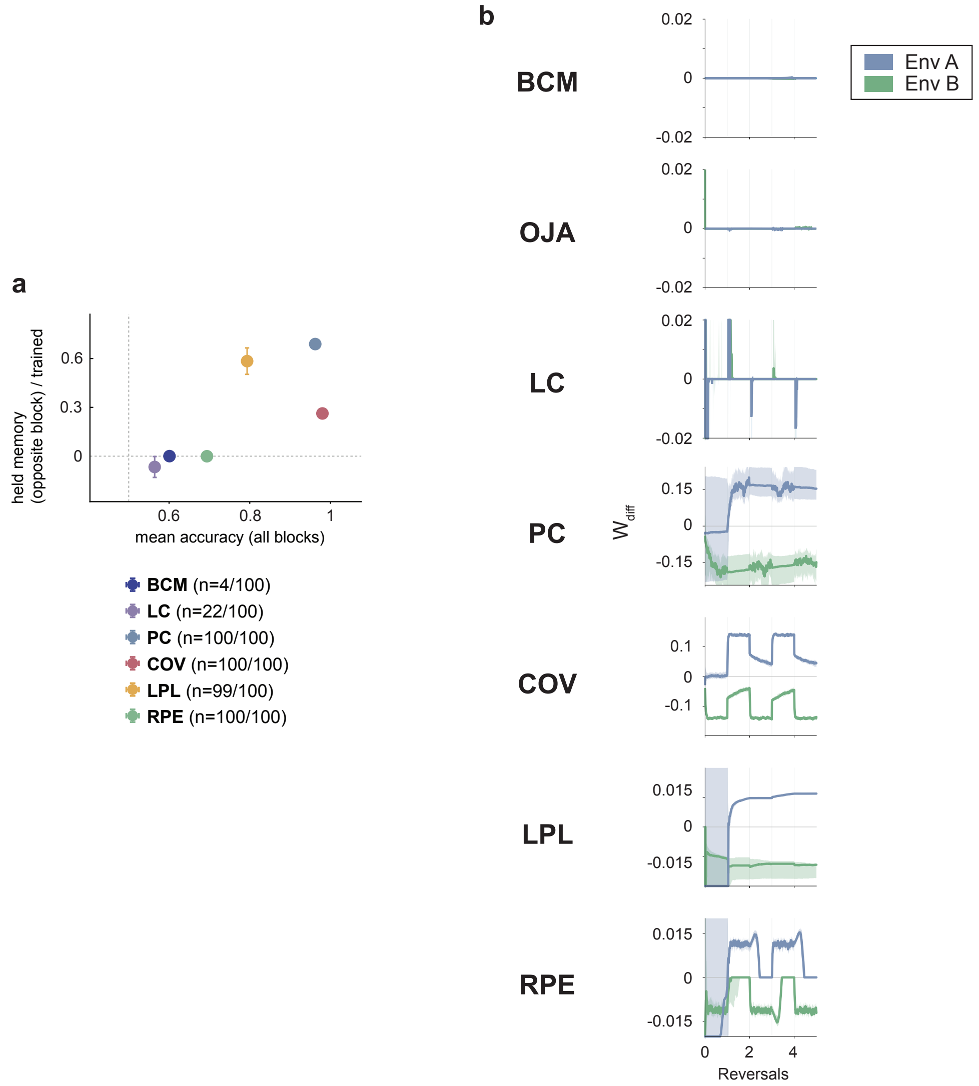


**Supplementary Figure S8: Memory retention across contexts: held memory versus accuracy.** **a**, Held memory during the opposite block vs. behavioral performance. Mean accuracy across all 10 reversal blocks ($x$) vs. retained memory of the idle context, as a fraction of each rule's own trained weight scale ($y$). Mean $\pm$ s.e.m. across seeds, coloured by rule. The top-right corner is best: it means that a rule both relearns the active context and keeps the memory of the one it just left (Section S3). OJA is not shown because none of its seeds had a usable trained scale (0/100); the BCM point is from its 4 usable seeds.

**b**, Weight differences ($W_{\text{diff}}$, from each context's input neuron) for each context and each rule across the first 5 of 10 reversal blocks ($n = 100$). Lines are medians and shading is the 35th–65th percentile range.

### Supplementary Methods

#### S1 Parameter selection

**Plasticity-off control.** To check that reward seeking at each operating point came from plasticity, we re-ran every operating point with $\beta = 0$ (50 seeds). With plasticity off, all rule–architecture pairs except BCM and LC in the large network scored 0.56–0.60, against 0.66–1.00 at their operating points. For BCM and LC in the large network, the unrestricted best cell ($\alpha = 1$, $\beta = 10^{-5}$) scored 1.00 with plasticity off. At $\alpha = 1$ each activity step overshoots its target and is clipped, so both motor neurons change by the same amount and their difference persists until an unrewarded choice drives activity to zero. This gives win–stay behavior without learning. We therefore excluded $\alpha = 1$ and $\beta = 10^{-5}$ for these two pairs. The selected cells (BCM: $\eta = 10^{-3}$, $\tau = 0.1$, $\alpha = 0.0848$, $\beta = 1$; LC: $\eta = 10^{-4}$, $\tau = 0.1$, $\alpha = 0.0072$, $\beta = 0.0848$; Table S2) score 0.68 and 0.74, against 0.58 and 0.56 with plasticity off. These two settings seek the rewarded arm only at large input magnitudes: at the end of 100,000 steps, the median P(correct) is below 0.5 for $r \leq 0.8$ (BCM) and $r \leq 0.6$ (LC), so neither has a measurable learning rate in Fig. 2c. OJA in the small network also sits at $\alpha = 1$ but scores above its plasticity-off control (0.66 vs. 0.58), so we kept it.

**Tie-breaking for the forgetting task.** For PC, RPE, COV and LPL, many cells reach the maximal deterministic-bandit score (26, 134, 362 and 370 cells, respectively). Among tied cells the argmax falls on the lowest admissible noise ($\eta = 10^{-5}$). With so little noise nothing perturbs the network away from its trained state, so "constant input" and "no input" become indistinguishable, and the mechanism under test is silenced. We therefore screened every tied cell (20 seeds; 100,000 training and 100,000 retention steps) and computed retention ratios as the mean $W_{\text{diff}}$ at the end of retention divided by the mean trained $W_{\text{diff}}$ (a ratio of means, robust to seeds with near-zero trained weights). We discarded cells with either ratio outside $(-0.3, 1.3)$ and selected the cell with the largest contrast between no-input and constant-input retention. For LPL the maximal-contrast cell proved unstable at full fidelity, so we used the tied cell closest to the ideal (retention 1, decay 0). OJA, BCM and LC have at most two tied cells and use their operating points. The resulting parameters (Table S2) are also used for the memory-retention task and, for PC, for the run-length analyses.

**Reward-magnitude mechanism (Fig. 2e–i).** The small-network PC operating point ($\eta = 10^{-5}$) is frozen: P(correct) stays near 0.57 and changes by less than 0.01 across reward magnitudes. We therefore chose $(\alpha, \beta) = (0.0848, 0.0164)$, one of the top-scoring cells at fixed $\eta = 10^{-2}$, the noise level of the large-network PC operating point. It has a separation of timescales between weights and activities ($\beta/\alpha \approx 0.19$). At $\eta = 10^{-2}$ the three-neuron network's asymptote still depends on reward (0.94 at $r = 0.2$ and 0.97 at $r = 1$ after 300,000 steps), because the accumulated activity noise is comparable to the output difference a learned weight produces at small $r$. We therefore lowered the noise to $\eta = 0.003$, at which the asymptote is 0.98–0.99 at every reward magnitude (Li & Erlich[5]).

#### S2 Reward-magnitude sweep

**Cells.** A cell was included if it chose the rewarded arm on more than 55% of all stored deterministic-bandit steps (averaged over seeds). This gave between 67 (OJA, small) and 2,613 (LPL, small) cells per rule and architecture, 12,880 in total. Every included cell was simulated at $r \in \{0.2, 0.4, 0.6, 0.8, 1.0\}$ with 20 seeds (base seed 235), with activities capped at 3 after the noise step.

**Budget.** All cells were first run for 20,000 steps. Cells whose median learning curve was still rising and below criterion at that budget were re-run for 100,000 steps, and their asymptotes were taken from the longer run.

**Metrics.** Rate and asymptote sensitivities were computed as in Methods (*Rate and asymptote sensitivity*), from 20 seeds per cell. Cells with fewer than three usable reward levels were excluded (9,279 cells retained).

#### S3 Memory-retention summary

For each seed, held memory is the weaker of the two channels' preferences while the other context is active. $\pi_A$ is averaged over the ends of B blocks, and $\pi_B$ over the ends of A blocks after the first. The minimum of the two is clipped to $[-0.3, 0.3]$ and divided by the seed's trained scale (Methods). Taking the minimum rather than the mean ensures that a rule tracking only the current contingency, which holds one association at a time, scores near zero. Accuracy is the fraction of correct choices in the second half of each block, averaged over the 10 blocks. Both are shown as mean ± s.e.m. across seeds with a usable trained scale.

### Supplementary Tables

**Supplementary Table S1: Statistics for the retention and forgetting comparisons (Fig. 5).** Paired one-sided Wilcoxon signed-rank test (no input vs. constant input) and one-sample one-sided Wilcoxon signed-rank test against 0 (reversal), on the tail-mean $W_{\text{diff}} = w_a - w_b$ ($a$, the originally rewarded arm) over the last 20% of the retention window; $n = 250$ seeds per rule; Benjamini–Hochberg FDR correction across 7 rules $\times$ 2 tests. No-input retention was significantly higher than constant-input retention in BCM, OJA, PC, COV, LPL and RPE (all $p_{\text{FDR}} < 10^{-8}$) but not LC ($p_{\text{FDR}} = 1.0$, wrong-signed). Reversal reached significantly negative $W_{\text{diff}}$ in PC, COV and RPE (all $p_{\text{FDR}} < 10^{-9}$) but not BCM, OJA, LC or LPL ($p_{\text{FDR}} = 1.0$). $r$, matched-pairs rank-biserial effect size ($+1$, every seed agrees with the tested direction; 0, no consistent direction). $^{***}p_{\text{FDR}} < 0.001$; n.s., not significant. $p_{\text{FDR}}$ values are floored at $10^{-9}$ for readability. Median retention as a fraction of each seed's trained $W_{\text{diff}}$ (no input vs. constant input): PC 0.81 vs. 0.34, COV 0.29 vs. 0.07, LPL 1.15 vs. 0.19, RPE 0.92 vs. 0.00. BCM and OJA stored almost no trained $W_{\text{diff}}$ (median $\approx 10^{-4}$), so their significant differences reflect constant input driving $W_{\text{diff}}$ negative (BCM median $-4.8$), not retention.

| | No input vs. constant input | | | Reversal vs. 0 | | |
|---|---|---|---|---|---|---|
| Rule | $r$ | $p_{\text{FDR}}$ | Sig. | $r$ | $p_{\text{FDR}}$ | Sig. |
| BCM | $+1.00$ | $< 10^{-9}$ | *** | $+0.97$ | 1.0 | n.s. |
| OJA | $+0.43$ | $2.7 \times 10^{-9}$ | *** | $+0.59$ | 1.0 | n.s. |
| LC | $-0.64$ | 1.0 | n.s. | $+1.00$ | 1.0 | n.s. |
| PC | $+0.41$ | $1.9 \times 10^{-8}$ | *** | $-1.00$ | $< 10^{-9}$ | *** |
| COV | $+0.98$ | $< 10^{-9}$ | *** | $-1.00$ | $< 10^{-9}$ | *** |
| LPL | $+0.97$ | $< 10^{-9}$ | *** | $+1.00$ | 1.0 | n.s. |
| RPE | $+1.00$ | $< 10^{-9}$ | *** | $-1.00$ | $< 10^{-9}$ | *** |

**Supplementary Table S2: Parameters by figure.** All simulations use $\kappa = 1$ and no activity cap unless listed. Arch.: S, small (1–2); L, large (1–50–2); CL, 2–2. "—" denotes rules without $\tau$.

**Reward-seeking operating points** (Fig. 1e–g, Fig. 2c)

| Rule | Arch. | $\alpha$ | $\beta$ | $\eta$ | $\tau$ |
|---|---|---|---|---|---|
| BCM | S | 0.0373 | 1 | $10^{-5}$ | $10^{-3}$ |
| BCM | L | 0.0848 | 1 | $10^{-3}$ | 0.1 |
| OJA | S | 1 | 1 | $10^{-5}$ | — |
| OJA | L | 0.00139 | 0.193 | $10^{-5}$ | — |
| LC | S | 0.000611 | 1 | $10^{-4}$ | 0.00464 |
| LC | L | 0.00720 | 0.0848 | $10^{-4}$ | 0.1 |
| PC | S | 0.0848 | 0.0373 | $10^{-5}$ | — |
| PC | L | 0.0164 | 0.00720 | $10^{-2}$ | — |
| COV | S | 0.000118 | 0.00316 | $10^{-5}$ | $10^{-3}$ |
| COV | L | 0.000611 | 0.00139 | $10^{-5}$ | $10^{-3}$ |
| LPL | S | 0.000118 | 0.000268 | $10^{-5}$ | $10^{-3}$ |
| LPL | L | 0.000268 | 0.000611 | $10^{-5}$ | $10^{-3}$ |
| RPE | S | $5.18 \times 10^{-5}$ | 0.0848 | $10^{-5}$ | — |
| RPE | L | $5.18 \times 10^{-5}$ | 0.193 | $10^{-5}$ | — |

**Forgetting and memory-retention parameters** (Fig. 5; Fig. S7 and Fig. S8; S and CL networks)

| Rule | $\alpha$ | $\beta$ | $\eta$ | $\tau$ | Selection |
|---|---|---|---|---|---|
| BCM | 0.0373 | 1 | $10^{-5}$ | $10^{-3}$ | operating point |
| OJA | 1 | 1 | $10^{-5}$ | — | operating point |
| LC | 0.000611 | 1 | $10^{-4}$ | 0.00464 | operating point |
| PC | 0.0373 | 0.00720 | $10^{-2}$ | — | plateau search (26 cells) |
| COV | 0.0164 | 0.0848 | $10^{-2}$ | 0.1 | plateau search (362) |
| LPL | 0.439 | 0.00720 | $10^{-2}$ | 0.00464 | plateau search, closest to ideal (370) |
| RPE | 0.439 | 1 | $10^{-2}$ | — | plateau search (134) |

**Explore–exploit representative cells** (Fig. 4b,c; Fig. S6; S network)

| Rule | $\alpha$ | $\beta$ | $\eta$ | $\tau$ |
|---|---|---|---|---|
| LC | 1 | 0.193 | $10^{-2}$ | 0.0215 |
| PC | 0.439 | 1 | $10^{-2}$ | — |
| COV | 0.000611 | 1 | $10^{-3}$ | 0.1 |
| LPL | 0.193 | 0.00316 | $10^{-3}$ | 0.0215 |
| RPE | 0.00139 | 0.00720 | $10^{-2}$ | — |

**Matching-pennies cells** (Fig. 3h–k; S network; selection in Methods)

| Rule | $\alpha$ | $\beta$ | $\eta$ | $\tau$ |
|---|---|---|---|---|
| BCM | 0.0373 | 1 | $10^{-4}$ | $10^{-3}$ |
| OJA | 0.00139 | 0.00139 | $10^{-2}$ | — |
| LC | 0.439 | 0.0848 | $10^{-6}$ | 0.0215 |
| PC | 0.0164 | 0.439 | $10^{-3}$ | — |
| COV | 0.00316 | 0.0373 | $10^{-6}$ | $10^{-3}$ |
| LPL | 1 | 1 | 0.1 | 0.1 |
| RPE | 0.0848 | 1 | $10^{-6}$ | — |

**Other panels** (PC, S network)

| Panel | $\alpha$ | $\beta$ | $\eta$ | Other | Seeds × steps |
|---|---|---|---|---|---|
| Fig. 1b,c | 0.01 | 0.01 | 0.005 | $s$: 1 then 0 | $1 \times 1{,}000$ |
| Fig. 2d | L operating point | | | $r = 0.1$–$1.0$ | 20 batches × 20 seeds × 20,000 |
| Fig. 3d | 0.023 | 0.15 | 0.005 | unbaited, 60 blocks, mean 5,000 trials | 20 |
| Fig. 4d | 0.0373 | 0.00720 | 0.01 | baited; ITI 0/2; $\kappa = 0.25$; cap 3 | $24 \times 120{,}000$ |
| Fig. 1j (stochastic) | 0.02 | 0.03 | 0.008 | deterministic bandit; cap 3 | $3 \times 40{,}000$ |
| Fig. 2e–i | 0.0848 | 0.0164 | 0.003 | $r \in \{0.2, \ldots, 1\}$; no cap | 800 (curves); 200 (drift) × 300,000 |
| Fig. 5c | 0.0373 | 0.00720 | 0.01 | retention conditions | $120 \times (100{,}000 + 20{,}000)$ |
| Fig. 3e–g | 0.01 | 0.02–0.15 | 0.005 | baited, total rate 0.45; cap 3 | $40 \times 40{,}000$ |
| Fig. 4e–h | 0.0373 | 0.00720 | 0.002 | baited 0.45/0.15; $\kappa = 0.25$; cap 3 | $1 \times 360{,}000$ per ITI |
| Fig. 4i–k | 0.1 | 0.01 | 0.005 | 5 blocks; $\kappa = 0.25$; cap 3 | $48 \times 100{,}000$ |
